# Many long intergenic non-coding RNAs distally regulate mRNA gene expression levels

**DOI:** 10.1101/044719

**Authors:** Ian C. McDowell, Athma A. Pai, Cong Guo, Christopher M. Vockley, Christopher D. Brown, Timothy E. Reddy, Barbara E. Engelhardt

## Abstract

Long intergenic non-coding RNAs (lincRNA) are members of a class of non-protein-coding RNA transcript that has recently been shown to contribute to gene regulatory processes and disease etiology. It has been hypothesized that lincRNAs influence disease risk through the regulation of mRNA transcription [88], possibly by interacting with regulatory proteins such as chromatin-modifying complexes [37, 50]. The hypothesis of the regulation of mRNA by lincRNAs is based on a small number of specific lincRNAs analyses; the cellular roles of lincRNAs regulation have not been catalogued genome-wide. Relative to mRNAs, lincRNAs tend to be expressed at lower levels and in more tissue-specific patterns, making genome-wide studies of their regulatory capabilities difficult [15]. Here we develop a method for Mendelian randomization leveraging expression quantitative trait loci (eQTLs) that regulate the expression levels of lincRNAs (linc-eQTLs) to perform such a study across four primary tissues. We find that linc-eQTLs are largely similar to protein-coding eQTLs (pc-eQTLs) in cis-regulatory element enrichment, which supports the hypothesis that lincRNAs are regulated by the same transcriptional machinery as protein-coding RNAs [15, 80] and validates our linc-eQTLs. We catalog 74 lincRNAs with linc-eQTLs that are in linkage disequilibrium with TASs and are in protein-coding gene deserts; the putative lincRNA-regulated traits are highly enriched for adipose-related traits relative to mRNA-regulated traits.

## 1 Introduction

Long intergenic non-coding RNAs (lincRNAs) are members of a class of non-protein coding RNA transcript greater than 200 nucleotides in length that do not overlap protein-coding gene annotations. LincRNAs have been shown to be necessary for various cellular and organismal processes, including mammalian X-chromosome inactivation [106], telomere maintenance [7], and cellular differentiation and specification [34, 51, 91, 95]. The dysregulation of lincRNAs has been noted in several important human phenotypes, including muscle performance, Alzheimer’s disease, and autoimmune diseases [2, 102]. A role for lincRNAs in disease is consistent with sequence conservation analyses, showing that they are under purifying selection [80]. However, lincRNA genes are generally less evolutionarily conserved than protein-coding exons or messenger RNA (mRNA) untranslated regions [36]. LincRNAs are thought to be regulated by the same transcriptional and RNA processing machinery as protein-coding mRNAs [15, 80]. Overall, however, lincRNAs are often expressed at lower levels and in a more tissue-specific patterns relative to mRNA [15, 38].

Of the thousands of lincRNAs recently annotated across human tissues using high-throughput sequencing technologies [15, 23, 73], only a small fraction have been functionally characterized [85]. There is a variety of functional mechanisms by which they are thought to act [9]. Nevertheless, a common theme has emerged that many lincRNAs might function to regulate the transcription of protein-coding genes [72, 88] through either cis (e.g., *XIST* [18]), trans (e.g., *Fendrr* [34]), or a combination of cis and trans mechanisms (e.g., *lincRNA-p21* [25, 46]). The best studied example of a regulatory lincRNA, *Xist*, is a ∼ 17 Kb transcript expressed in Eutherian mammalian females from only one of two X-chromosomes [13]. Following capping, splicing, and polyadenylation, *Xist* bypasses nuclear export pathways to localize to the X-chromosome inactivation center, where it accumulates and spreads in cis [18]. *Xist* coats the X-chromosome and recruits the Polycomb Repressive Complex 2 (*PRC2*) [106], which promotes the formation of the repressive chromatin mark H3K27me3. Interactions with chromatin-modifying complexes have also been noted for many other lincRNAs [37, 50, 89, 97, 105], although there are other mechanisms by which lincRNAs can act to affect gene expression. In particular, lincRNAs have been experimentally shown to bind to transcription factors and alter their activity [39], stabilize mRNA transcripts [53], and possibly affect the transcription of neighboring mRNA genes by influencing local transcriptional environments [78]. However, regardless of detailed functional characterization of individual lincRNAs, there is no consensus on the extent to which lincRNAs regulate transcription or the modes by which they might act.

Here we aim to systematically characterize the global transcriptional effects of many lincRNAs simultaneously and the modes by which they regulate transcription. There are two primary statistical challenges to this work. First, correlations between genes do not discriminate between causal interactions, where the lincRNA directly regulates gene transcription, and unobserved regulatory factors, where the transcription of both the lincRNA and another gene is co-regulated. Second, studies on lincRNAs have reduced statistical power because lincRNAs are on average expressed at much lower levels than mRNAs, and lincRNA annotations are of lower quality than mRNA annotations. We are able to overcome both of these challenges by using associations between genotypes and RNA expression levels to resolve causal models and to control for statistical power differences between mRNAs and lincRNAs. Genetic variants that are associated with RNA expression, or expression quantitative trait loci (eQTLs), have been extensively mapped in cell lines and tissue types using genome-wide approaches [28, 32, 64, 70, 74, 79]. Recent eQTL studies that include lincRNAs have shown that genetic variation also contributes to the regulation of lincRNA expression levels [67, 81].

More broadly, non-coding variants are thought to affect organismal phenotypes largely through the modulation of gene expression levels [69]. Support for this comes from the finding that trait-associated SNPs (TASs) are enriched in collections of protein-coding expression quantitative trait loci (eQTLs) across human cell lines and tissues [71]. Genome-wide association studies (GWAS) have shown that the majority of TASs are non-coding. Some of these TASs are located in protein-coding gene deserts [62, 66] and in lincRNAs [15], such as *MIAT*, which is associated with myocardial infarction [47]. Other TASs include eQTLs for lincRNAs like *PTCSC3*, which is associated with papillary thyroid carcinoma [49]. However, as with transcription regulation, the role of lincRNAs in regulating complex traits is unclear on a genome-wide level.

To better understand the contribution of lincRNAs expressed in primary tissues on complex traits and mRNA expression levels, we performed a systematic eQTL study to identify genetic variants regulating lincRNA and mRNA expression levels across multiple tissues. We used these genetic associations in statistical enrichment models to study shared transcriptional mechanisms across lincRNAs and mRNAs via their cis-eQTLs. We used similar enrichment analyses to compare the roles of lincRNAs and mRNAs on the regulation of complex trait types. We used a genome-wide approach of Mendelian randomization to quantify the effects of lincRNAs on distal mRNA expression levels across the genome, and investigated several possible mechanisms of transcription regulation in the regulatory lincRNAs.

## 2 Results

We performed a genome-wide eQTL association study using RNA-sequencing data and array-based, imputed genotype data to identify genetic variants that influence lincRNA gene expression levels (*linc-eQTLs*) and mRNA (protein-coding) gene expression levels (*pc-eQTLs*). We used data from four primary tissues—adipose (*n* = 107 samples, 103 individuals), artery (*n* = 118 samples, 117 individuals), lung (*n* = 133 samples, 121 individuals), and skin from the lower leg (*n* = 109 samples, 105 individuals)— from 167 donors to the Genotype-Tissue Expression (GTEx) project pilot v4 data [6].

We adopted the definition of cis- and trans-regulation current in the eQTL literature as association between genotype at a locus and expression of a gene where these genomic elements are separated by less than or greater than some distance threshold (here, < 100 Kb and > 1 Mb), respectively. This definition is agnostic of functional mechanism, and is only a proxy for the true definition of cis-regulation, which is allele-specific regulation; nevertheless it is a useful proxy for the study of genome-wide patterns of transcriptional regulation by lincRNAs.

### 2.1 Cis-eQTL discovery

We tested for association between the genotype of single nucleotide polymorphisms (SNPs) and gene expression levels in each tissue separately; we also tested for association jointly across all tissues using a multi-tissue association test [30]. More specifically, we tested for association of a gene with cis-SNPs within the interval from 100 Kb upstream of the transcription start site (TSS) to 100 Kb downstream of the transcription end site (TES; Supplemental Table 1). We computed Bayes factor (BF) thresholds corresponding to FDR ≤ 5% for both methods separately using permutations and took the union of significant SNP-gene associations to produce the full set of eQTLs. We found 126,258 SNPs associated with 1,567 of the 4,368 lincRNA genes, or 36% of the lincRNA genes tested (FDR ≤ 5%; Supplemental Fig. 1, Supplemental Table 2). In contrast, we identified 905, 704 SNPs associated with 9, 794 of the 18, 191 protein-coding genes, or 54% of the mRNA genes tested (FDR≤ 5%; Supplemental Fig. 1, Supplemental Table 2). We selected the single most significant eQTL for each gene—where the single- and multi-tissue analyses differed, we included both of the most significant eQTLs—to define a set of *best eQTLs* that included 2,034 cis-linc-eQTLs and 13, 522 cis-pc-eQTLs. We found similar enrichment of the best cis-eQTLs proximal to the TSS and TES for both lincRNAs and mRNAs (Fig. 1A and B) [100]. Linc-eQTLs and pc-eQTLs were similarly overrepresented around annotated splice-sites, suggesting that lincRNA expression levels might be subject to the same degree of post-transcriptional regulation as mRNAs (Supplemental Fig. 2).

**Figure 1.**
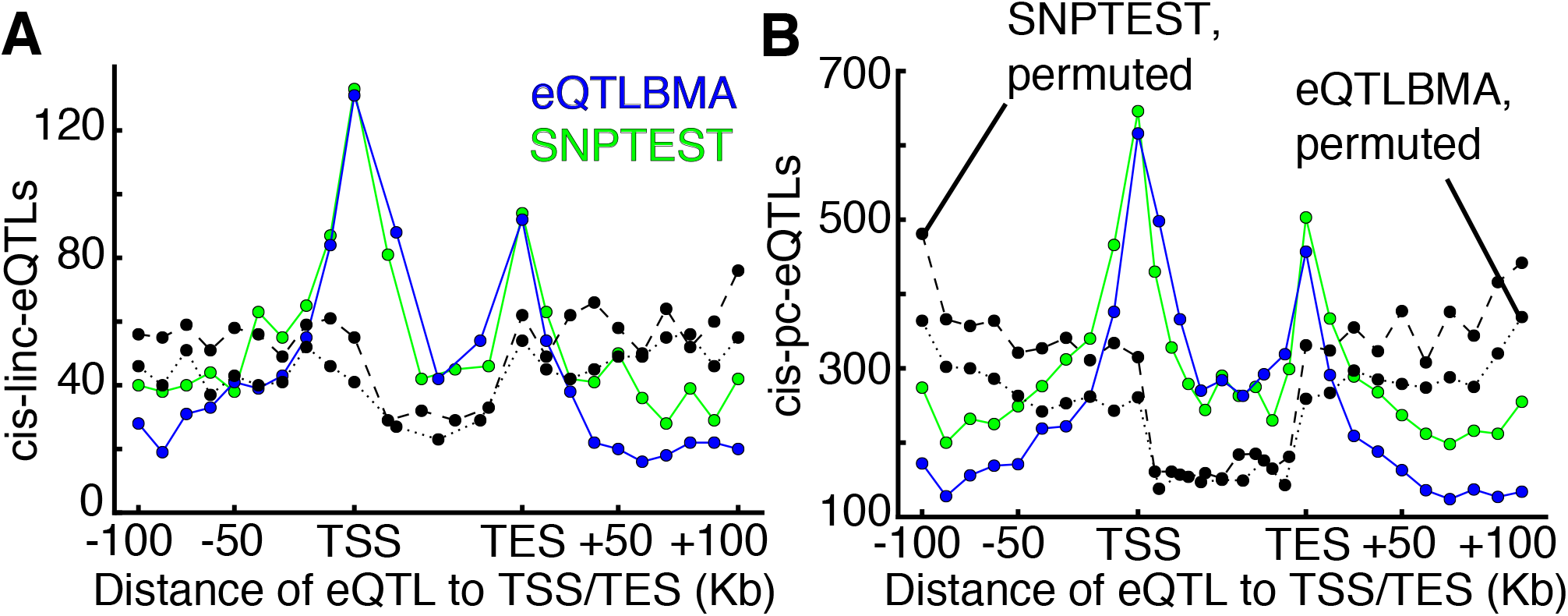
Linc-eQTL and pc-eQTL associations across four tissue types. Distribution of the location of the best cis-eQTL for the transcription start and end sites of each gene for **Panel A**, lincRNA and **Panel B**, mRNA.

We quantified the relative tissue specificity of linc-eQTLs, and we found that the number of lincRNAs with at least one linc-eQTL was consistent across the four tissues (Fig. 2A, B, and Supplemental Fig. 3A). Moreover, 35% of linc-eQTLs were unique to a single tissue (Fig. 2A). The number of mRNAs with at least one cis-pc-eQTL was also consistent across tissues (Fig. 2B and Supplemental Fig. 3B), and 31% of pc-eQTLs were unique to a single tissue. Controlling for gene length, GC content, number of exons, average expression levels, SNP minor allele frequency (MAF), and tissue-specificity of expression [15], linc-eQTLs were significantly more tissue-specific than pc-eQTLs (Eqn.1; Student’s t-test; *p* ≤ 5.1 × 10^−12^).

**Figure 2.**
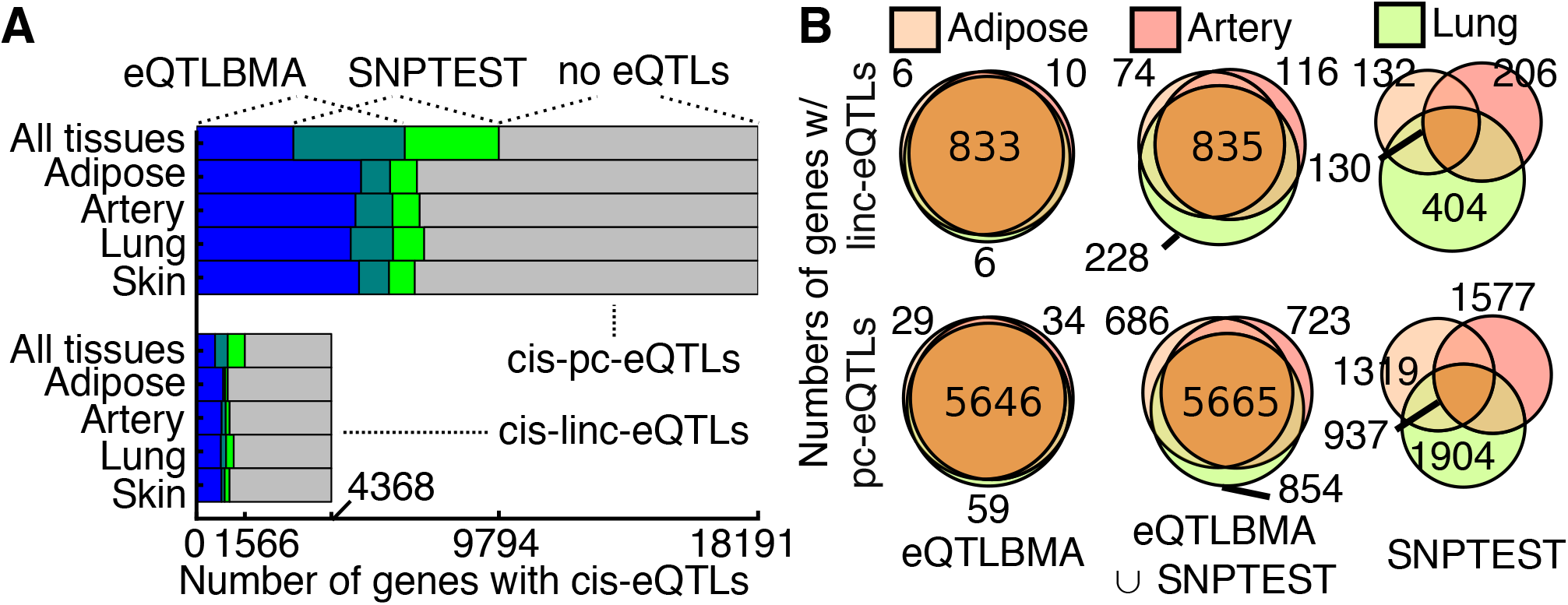
Genes with cis-eQTLs within and across tisuses. **Panel A**, Numbers of genes with cis-eQTLs in each tissue and across all tissues by discovery association method. **Panel B**, Numbers of genes with cis-eQTLs that are shared and unique across tissues (skin not shown, see Supplemental Fig. 2) by discovery association method.

We found that tissue-specific linc- and pc-eQTLs had lower median association significance than non-tissue-specific eQTLs [24] (Mann-Whitney U-test, linc- and pc-eQTLs, *p* ≤ 1×10^−100^). Small sample size in the pilot GTEx data, intrinsic tissue specificity of lincRNA expression, and poor lincRNA annotations may affect these results.

The majority of cis-linc-eQTLs and cis-pc-eQTLs were associated with only one gene; however, we found substantial overlap between SNPs identified as both cis-linc-eQTLs and cis-pc-eQTLs. Specifically, 27, 524 SNPs were significantly associated with expression levels of both a lincRNA and an mRNA. Shared cis-eQTLs were 2.3 times more likely to have the same direction of effect on lincRNA and mRNA expression levels, rather than opposite direction of effect (Fisher’s exact test, *p* ≤ 1×10^−100^; Supplemental Fig. 4). We did not find any overrepresented patterns of the shared eQTL with respect to the transcription orientation of the lincRNA and mRNA in a genomic locus across matched and unmatched effects. Using conditional analyses, we found that the large majority of eQTLs that were significant cis-linc-eQTLs and cis-pc-eQTLs were regulating both gene targets directly (Eqn. 2, Supplemental Fig. 5A). This scenario also allows for a common but unmodeled effect (e.g., a shared transcription factor that regulates both the mRNA and the lincRNA). For a small minority of shared SNPs, we identified a direct and an indirect effect. These results agree with the lack of significant patterns of orientation of the mRNA and lincRNA gene bodies with respect to the shared cis-eQTL. These results suggest that lincRNA and mRNA transcription share similar local non-genic mechanisms of regulation. They also suggest that there do not exist ubiquitous patterns of local mRNA-lincRNA regulatory interactions, although it is possible that this lack of signal is due to insufficiently powerful statistical models to resolve these local interactions.

We considered the possibility that a SNP-gene association may be an artifact of the RNA-seq platform or read mapping pipeline by examining these eQTLs in nine publicly available eQTL data sets from seven different (unmatched) cell types [12]. In particular, we looked for each SNP-gene pair in the association mapping results from each study. We removed all genes not assayed on the gene expression array from the data and all SNPs that were not tested for association in that study. We considered the association to replicate in a study when the gene-SNP pair had a log_10_ *BF* ≥ 1. Overall, 44 cis-linc-eQTLs were tested; of these, 20 linc-eQTLs replicated (45%). Similarly, 7, 989 cis-pc-eQTLs were tested; of these, 4, 361 pc-eQTLs replicated (55%). Although the number of potential associations tested differs substantially, linc-eQTLs and pc-eQTLs replicated across platforms and cohorts at similar frequencies (Fisher’s exact test, *p* ≤ 0.6), indicating that well-annotated lincRNAs and mRNAs are similarly reproducible across RNA-seq and microarray platforms.

### 2.2 Similar cis-regulatory element enrichment for linc-eQTLs and pc-eQTLs

Many of the mechanisms by which cis-eQTLs regulate transcription of mRNAs are well understood [32]; to validate our cis-linc-eQTLs, we compared enriched co-localization of cis-linc-eQTLs with specific classes of cis-regulatory elements (CREs) to enriched co-localization of cis-pc-eQTLs with these CREs [12]. We performed enrichment analyses of our cis-linc-eQTLs and cis-pc-eQTLs with genome-wide maps of several indicators of CREs—DNase I hypersensitive sites (DHSs), nine histone modifications, and binding of RNA Polymerase II, *EZH2*, and *CTCF*—identified in nearest match cell lines from the ENCODE project [20]. Then we modeled CRE-eQTL co-localization using logistic regression controlling for distance to TSS, average gene expression level, and MAF [12] (Eqn. 3) to statistically quantify enrichment.

The best cis-linc-eQTLs and the best cis-pc-eQTLs were significantly enriched in CREs with features indicative of active gene transcription (putatively *active CREs*), including DHSs (Fig. 3C), H3K27ac (Fig. 3B), H3K9ac, H3K4me3, and RNA Polymerase II binding sites (Supplemental Fig. 6). Among all CREs, cis-pc-eQTLs and cis-linc-eQTLs were most differentially enriched for overlap with H3K36me3 (Fisher’s exact test, best: *p* ≤ 2.2 × 10^−16^; Fig. 3C). A recent study found that protein-coding genes have elevated H3K36me3 signals downstream of the TSS across cell types, while lncRNA genes do not [90]. The differential enrichment of eQTLs in H3K36me3 sites may indicate differential usage of this histone modification in transcription regulation of protein-coding genes as compared with lincRNA genes. For the remainder of the CREs, however, the signal for enrichment in active CREs was similar for cis-linc-eQTLs and cis-pc-eQTLs, indicating similar allele-specific transcription regulatory mechanisms and globally validating our cis-linc-eQTLs.

**Figure 3.**
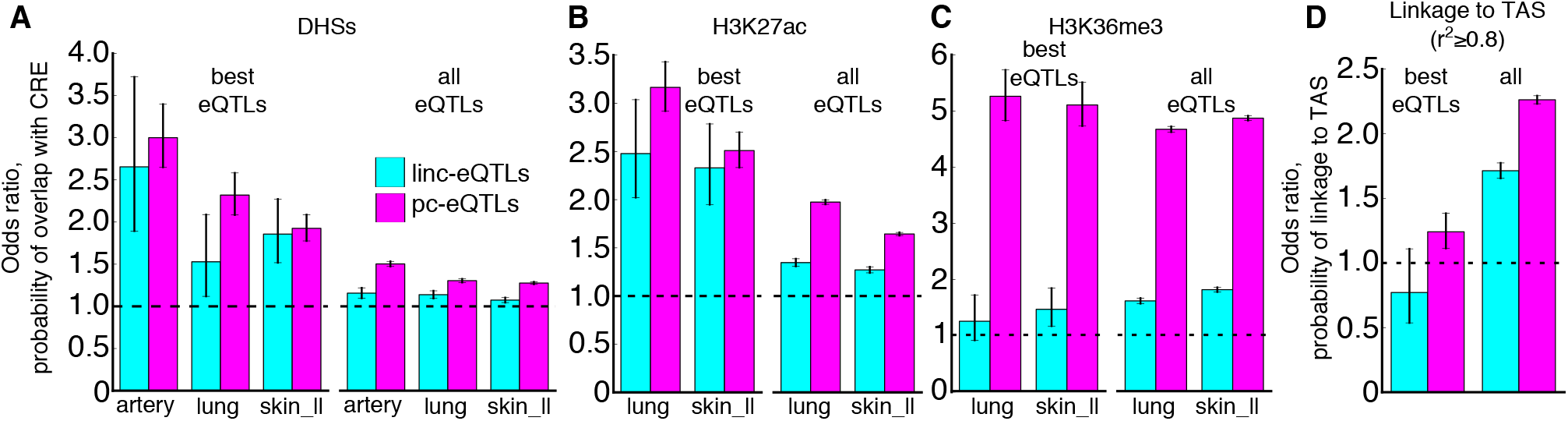
Enrichment of overlap with CREs and TASs for cis-linc-eQTLs and cis-pc-eQTLs. **Panel A**, Odds ratio of probability of overlap with DHSs from ENCODE for the best and all cis-linc-eQTLs and the best and all cis-pc-eQTLs across each tissue type with available matched ENCODE cell-type data. **Panel B**, Enrichment of eQTLs in histone modification H3K27ac. **Panel C**, Enrichment of eQTLs in histone modification H3K36me3. **Panel D**, Odds ratio of probability of linkage to a TAS for the best and all cis-linc-eQTLs and cis-pc-eQTLs.

### 2.3 Enrichment of cis-linc-eQTLs with trait associated SNPs

Previous studies have shown that pc-eQTLs are enriched in trait-associated SNPs (TASs), or SNPs identified via genome-wide association studies (GWAS) for organismal traits, suggesting that genetic regulation of mRNA transcription contributes to complex organismal traits [12, 71]. To study the impact of the genetic regulation of lincRNA transcription on organismal traits, we tested for enrichment of TASs from the NHGRI GWAS Catalog [43] among cis-linc-eQTLs and cis-pc-eQTLs. The probability of linkage to a TAS was modeled with logistic regression controlling for distance to TSS, average expression level, and SNP MAF, requiring the TAS and eQTL to have *r*^2^ ≥ 0.8 (Eqn. 4). Both cis-linc-eQTLs and cis-pc-eQTLs were enriched for linkage to TASs (Fig. 3D).

We compiled a set of 74 lincRNAs that had cis-linc-eQTLs in LD with TASs (*r*^2^ ≥ 0.8) that were not in LD (*r*^2^ < 0.2) with any cis-pc-eQTLs or local-pc-eQTLs (*local* indicates a distance of SNP to gene < 1 Mb) and were also at least 10 Kb from the nearest protein-coding gene, suggesting that the trait may be regulated through the lincRNA (*explanatory lincRNAs*; Supplemental Table 3). Eighty-eight TASs associated with a total of 96 traits were explained by these 74 explanatory lincRNAs (Supplemental Table 4). Among the several trait types enriched among TASs linked to linc-eQTLs but not to pc-eQTLs, *obesity-related traits* had the greatest enrichment, making up 8% of the 96 traits (Fisher’s exact test, *p* ≤ 0.029).

To explore the possible regulatory mechanisms of our explanatory lincRNAs, we tested these lincRNAs for possible mechanistic signatures (see Methods), including secondary structure conservation and protein-coding potential. None of these signatures were found in notable proportions of the explanatory lincRNAs, consistent with the notion that the functional mechanisms of lincRNAs are highly variable [98]. For all mechanistic analyses, we found that explanatory lincRNAs were not able to be distinguished from non-explanatory lincRNAs with cis-linc-eQTLs after controlling for transcript length. Two of our explanatory lincRNAs are likely to code for protein (*LINC00452* and *PCAT1*; Supplemental Table 3), but this does not appear to be common despite recent work using ribosomal profiling [10] and functional characterization [3]. The variety of possible mechanisms and the lack of evolutionary conservation of lincRNAs between humans and model organisms, makes experimental characterization of a single explanatory lincRNA difficult.

Three of the TASs that we tested here were previously found to be associated with traits related to adipose tissue (*adiposity* [56] and *visceral adipose tissue/subcutaneous adipose tissue ratio* [31]). These three TASs were located in a single genomic locus in strong LD with 132 cis-linc-eQTLs for the explanatory lincRNA *RP11-392O17*, with no cis- or local-pc-eQTLs in LD with the TAS (Fig. 4). Eight additional TASs are in LD (*r*^2^ ≥ 0.8) with cis-linc-eQTLs for this same lincRNA in this gene desert, seven of which were adipose-related traits (adiponectin levels [22], fasting insulin-related traits [61], visceral adipose tissue/subcutaneous adipose tissue ratio [31], hip-adjusted BMI [86], and waist-hip ratio [41]) and the non-adipose related trait was a sexually dimorphic trait (osteoarthritis [5]); although these eight additional TASs were also in weak LD (0.2 < *r*^2^ < 0.8) with a singleton cis-pc-eQTL in skin for gene SLC30A10, the explanatory value of this association is weak relative to the explanatory lincRNA. The cis-linc-eQTL association was most significant in adipose tissue. One of these linked cis-linc-eQTLs, rs2605100, has been found to be associated with adiposity [56] and resides over 200 Kb away from the nearest protein-coding gene, *LYPLAL1*. Furthermore, *LYPLAL1* does not have a local-pc-eQTL in LD with any of the adipose-related TASs. Our data show that, of the four tissues in this study, lincRNA *RP11-392O17* is most highly expressed in adipose tissue (Fig. 4); the only tissue in which this lincRNA has higher levels of expression is pancreas, which produces a number of sugar-metabolizing hormones, including insulin and glucagon. In aggregate, these results suggest that the cis-linc-eQTLs associated with this lincRNA may be involved in regulating risk of this collection of adipose-related traits.

**Figure 4.**
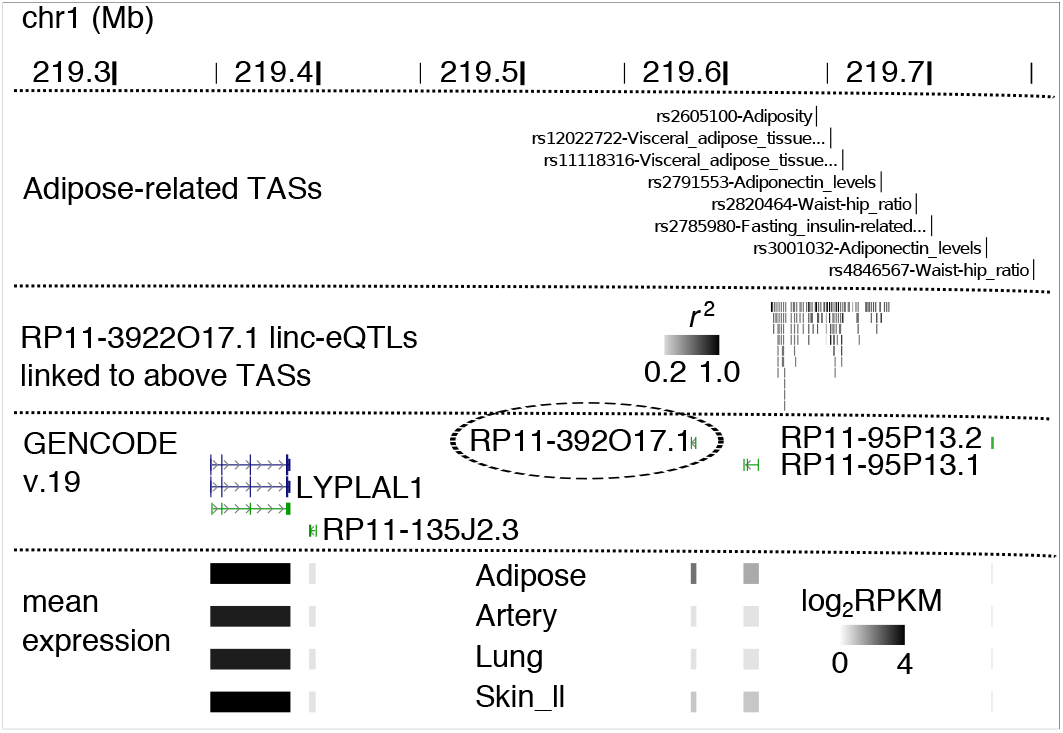
Genomic environment of ten adipose-related TASs. The genomic environment around ten adipose-related TASs and cis-linc-eQTLs for the lincRNA *RP11-392O17.1.* Cis-linc-eQTLs linked to at least one adipose-related TAS with *r*^2^ > 0.8 are shown. Gene models from GENCODE v. 19 shown for all genes in window (lncRNA *RP11-135J2.4*, which overlaps *LYPLAL1*, omitted). Mean log_2_ (*RPKM*) per tissue per gene is shown proportional to the highest expression estimate in the genomic window (*LYPLAL1* in adipose tissue).

We mention five other explanatory lincRNAs that are good candidates to mediate disease risk, based on their long distance from the nearest protein coding gene, the number of linked TASs, and the effect size of the cis-linc-eQTLs. First, lincRNAs *RP11-815M8.1* and *RP11-400N13.2* both were explanatory lincRNAs for TASs for progressive supranuclear palsy [44] and colorectal cancer [45] (Supplemental Table 3, 4). Second, lincRNA *RP11-400N13.1* had cis-linc-eQTLs that were strongly linked to one TAS for keloid (scar formation) [68]. Third, lincRNA *RP11-345M22.2* was strongly linked to four TASs for urate levels [52], thyroid volume [96], thyroid hormone levels [82], and thyroid function [87] (Supplemental Table 3, 4). Fourth, *PCAT1* was strongly linked to two obesity-related trait SNPs [19], but based on our analyses appears to be protein coding (Supplemental Table 3). Experimental characterization of these disease-related lincRNAs is difficult as described earlier due to low sequence conservation and other factors; moreover, none of these five lincRNAs appear to be small peptide hormones. However, our work indicates that including cis-linc-eQTLs in functional analyses of TASs is essential, as certain organismal traits may be regulated through lincRNAs.

Our results on the enrichment for linkage to TASs among cis-eQTLs defined linkage between an eQTL and a TAS if the two SNPs were located within 100 Kb and had a pairwise *r*^2^ ≥ 0.8, yet our results were robust to different lower bounds on *r*^2^ between the TAS and the cis-eQTL. Specifically the *all linc-eQTL* and *all pc-eQTL* sets were enriched for linkage to TASs at *r*^2^ ≥ {0.20-0.95}. The *best pc-eQTL* set was enriched for linkage to TASs at *r*^2^ ≥ {0.20-0.90}. The *best linc-eQTL* set was not enriched for linkage to TASs at any tested *r*^2^ threshold.

### 2.4 Mechanistic hypotheses for candidate lincRNAs

We found that linc-TASs were significantly enriched for *obesity-related traits* compared to pc-TASs. This enrichment generally increased with increasing stringency in defining explanatory linc-eQTLs by distance to the nearest protein-coding gene—and conversely defining explanatory pc-eQTLs by distance to the nearest lincRNA gene (Fisher’s exact test, *r*^2^ > 0.2: *p* ≤ 0.007; *r*^2^ > 0.8: *p* ≤ 0.035; *r*^2^ > 0.2 and > 10 Kb constraint: *p* ≤ 2.1 × 10^−4^; *r*^2^ > 0.8 and > 10 Kb constraint: *p* ≤ 0.028; *r*^2^ > 0.2 and > 100 Kb constraint: *p* ≤ 2.1 × 10^−5^; *r*^2^ > 0.8 and > 100 Kb constraint: *p* ≤ 0.01). We used thresholds of *r*^2^ > 0.8 to define eQTL-TAS linkage, *r*^2^ < 0.2 to characterize no eQTL-TAS linkage, and the protein coding gene > 10 Kb away from the TAS. We found that six lincRNAs had linc-eQTLs linked to TASs for obesity-related traits that were not linked to pc-eQTLs or nearby protein coding genes, including **PCAT1** and *PWRN1*, known to be functionally important in prostate cancer [83] and Prader-Willi syndrome [14], respectively.

Currently, few lincRNAs have experimentally verified functions or disease associations. Of these, only *PCAT1* from the Long Non-Coding RNA Database (lncRNAdb v.2.0) [85] was found in our candidate set. In terms of coding potential, the candidate set could not be distinguished from the set of its reverse complement sequences. We tested for coding potential, evolutionary conservation, conserved secondary structure, secondary structure folding energy, number of miRNA binding sites, nuclear versus cytoplasm localization, and signal peptide potential; for none of these tests could the candidate lincRNAs be distinguished from their complement set among all lincRNAs tested for cis-linc-eQTLs after controlling for transcript length. This finding is consistent with the notion that lincRNAs are a heterogeneous group of molecules whose functional mechanisms vary substantially [98].

Several of our candidate lincRNAs are likely protein-coding genes according to *CPAT, LINC00452* and **PCAT1** in particular (Supplemental Table 4). Additional work is needed to describe the potential action of other promising explanatory lincRNAs, such as *RP11-815M8.1* and *RP11-345M22.2*, which do not have a clear mechanism based on these analyses.

### 2.5 Experimental validation of linc-eQTLs

To experimentally confirm the regulatory effects of cis-linc-eQTLs on transcription, we selected seven loci that included cis-linc-eQTLs and cis-pc-eQTLs to validate using allele-specific luciferase reporter assays. We chose cis-eQTLS that were located in DHS shared across cell types that represent the tissues used, and compared relative enhancer activity between the major and minor haplotype. Of the seven loci tested, four were cis-pc-eQTLs, and four were cis-linc-eQTLs (one was both a cis-pc-eQTL and a cis-linc-eQTL). Three of the four cis-linc-eQTLs had a significant difference in luciferase expression between the major and minor haplotypes and in all cases the direction of the effect agreed with the association analysis (Student’s t-test; *p* ≤ 0.05, Student’s t-test; Fig. 5). Of the four cis-pc-eQTLs, three had a significant effect on luciferase expression. Two of those agreed in direction with the association analysis, and one acted in the opposite direction. The latter cis-pc-eQTL tested a tissue-specific association, potentially explaining the single inconsistent result. Overall, these results serve both as positive validation for the association analysis, and also support the hypothesis that genetic variation in DHSs are largely responsible for allele-specific transcription regulation.

**Figure 5.**
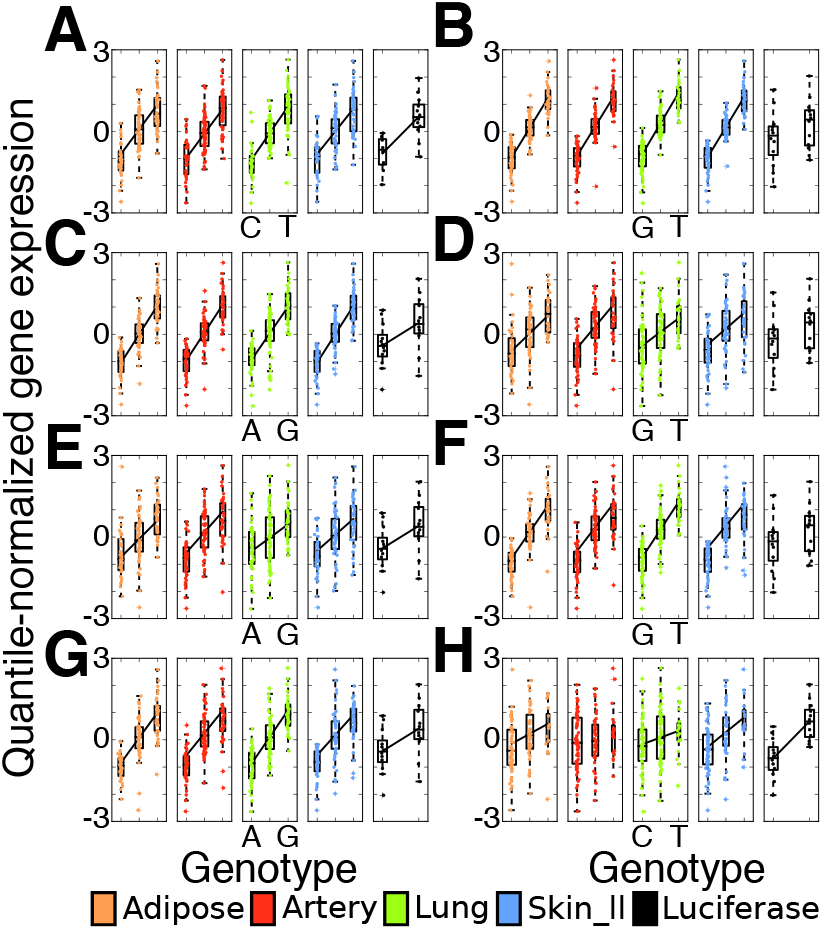
Experimental validation results for selected cis-linc-eQTLs. A regression line indicates an association between genotype and gene expression significant at the genome-wide level (tissues), or a significant association in the luciferase assay: **Panel A**, ENSG00000198468.3 versus rs1047881 (*p* ≤ 2.8 × 10^−3^), **Panel B**, ENSG00000204792.2 versus rs12366 (*p* > 0.05), **Panel C**, ENSG00000204792.2 versus rs13024157 (*p* ≤ 1.5 × 10^−4^), **Panel D**, ENSG00000230836.1 versus rs12366 (*p* > 0.05), **Panel E**, ENSG00000230836.1 versus rs13024157 (*p* ≤ 1.5 × 10^−4^), **Panel F**, ENSG00000236209.1 versus rs12366 (*p* > 0.05), **Panel G**, ENSG00000236209.1 versus rs13024157 (*p* ≤ 1.5 × 10^−4^), **Panel H**, ENSG00000269893.2 versus rs658642 (*p* ≤ 1.1 × 10^−2^).

**Figure 6.**
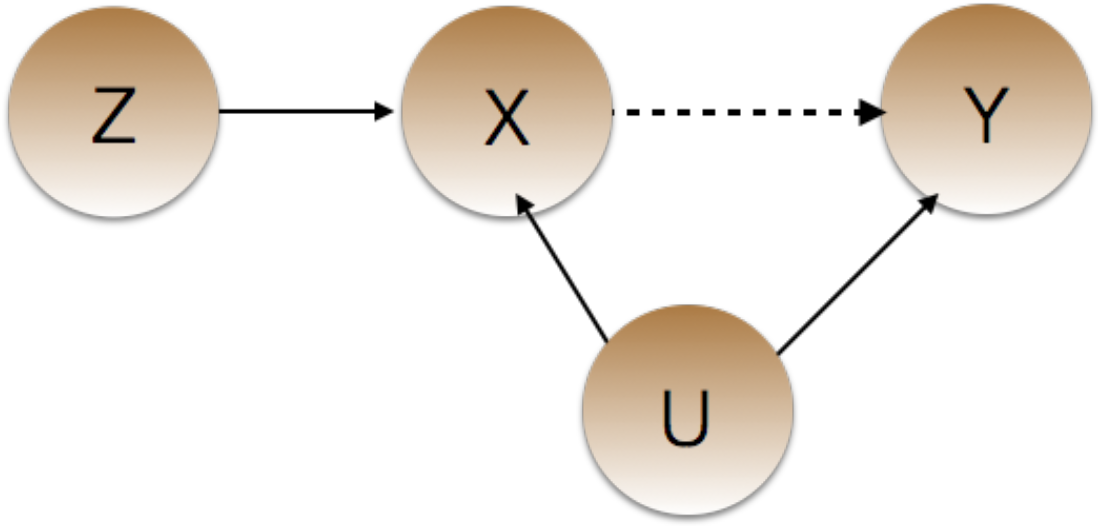
Cartoon capturing Mendelian randomization. Here *z* is the *instrumental variable* (i.e., a SNP) that directly affects the *independent variable x* (i.e., the cis-RNA), which, in turn, may or may not affect the *dependent variable y* (i.e., trans-mRNA). Both *x* and *y* may be affected by unmeasured variables u. Solid arrows represent observed relationships; the dashed line is the relationship that MR quantifies.

### 2.6 Mendelian randomization to test for RNA-mRNA regulation

The predominant hypothesis about the cellular role of lincRNAs is that they regulate the transcription of protein-coding genes. Previous studies have shown that i) lincRNAs interact with chromatin-modifying complexes [37, 50]; ii) lincRNA knockdown affects expression levels of protein-coding genes [37]; iii) lincRNAs, including *ROR, MALAT1 MEG3, MEG8, TERC*, and others, have been experimentally shown to function as trans-regulators [16, 17, 37, 57, 65, 107]. To do this, we tested for the trans-regulatory effects of lincRNAs in two ways: a n¨aive approach and an approach using a Mendelian randomization test.

#### 2.6.1 Näive approach for testing for trans-effects of lincRNAs

In our first näive approach, we tested the best cis linc- and pc-eQTLs for association with the expression levels of all genes with non-zero expression in at least 10% of samples in all four tissues in a cross-tissue manner. We designed this test for a general transcriptional role for lincRNAs using linc-cis-eQTLs for two reasons: first, using the genotypes instead of the lincRNA expression levels allowed us to control for potential loss of statistical power due to lower expression levels of lincRNAs versus mRNAs; second, we were able to calibrate our results against the same results from cis-pc-eQTLs, which had an similar distribution of effect sizes to the cis-linc-eQTLs and a substantial number of known transcription regulators. We did not find a significant difference between the distributions of the magnitude of effect sizes for the eQTLBMA best cis-linc-eQTLs and best cis-pc-eQTLs (Mann-Whitney U-test, *p* > 0.05; Supplemental Fig. 7A). However, the SNPTEST best cis-linc-eQTLs had slightly but significantly greater absolute effect sizes than cis-pc-eQTLs (Mann-Whitney U-test, *p* = 1.04 ×10^−4^; Supplemental Fig. 7B). If this slight difference in effect sizes affected results, it should advantage cis-linc-eQTLs over cis-pc-eQTLs. This approach let us statistically compare the proportion of lincRNAs having a trans effect on a protein coding gene to the proportion of protein coding genes having a trans effect on a protein coding gene; however, this approach does not control for shared latent confounders.

Using the näive approach, we found 9 cis-linc-eQTLs that were associated with 10 (0.05% of the total) protein-coding genes across four tissue types, resulting in 11 trans-pc-eQTLs (FDR ≤ 20%; Supplemental Table 2). For comparison, we found 7,594 cis-pc-eQTLs were associated with 7,046 (38.7%) protein-coding genes in trans across four tissue types, resulting in 22, 527 trans-pc-eQTLs (FDR ≤ 20%; Fig. 7A and Supplemental Table 2). All trans-pc-eQTLs were identified via multi-tissue mapping, while only 39 trans-pc-eQTLs were replicated using single-tissue mapping, pointing to the increased power of multi-tissue analyses in identifying trans-pc-eQTLs. Thus, using this näive approach, we observed a 2, 000 times greater chance that an mRNA will be associated in trans with a cis-pc-eQTL than with a cis-linc-eQTL. While 85% of cis-pc-eQTLs were estimated to be shared across all four tissues using eQTLBMA, only 11% of trans-pc-eQTLs (from cis-pc-eQTLs) were estimated to be shared across all four tissues. Our results provide support for the hypothesis that trans-eQTLs are more tissue-specific than cis-eQTLs [29, 35, 84]; however, we have limited power in these pilot data to resolve this question.

**Figure 7.**
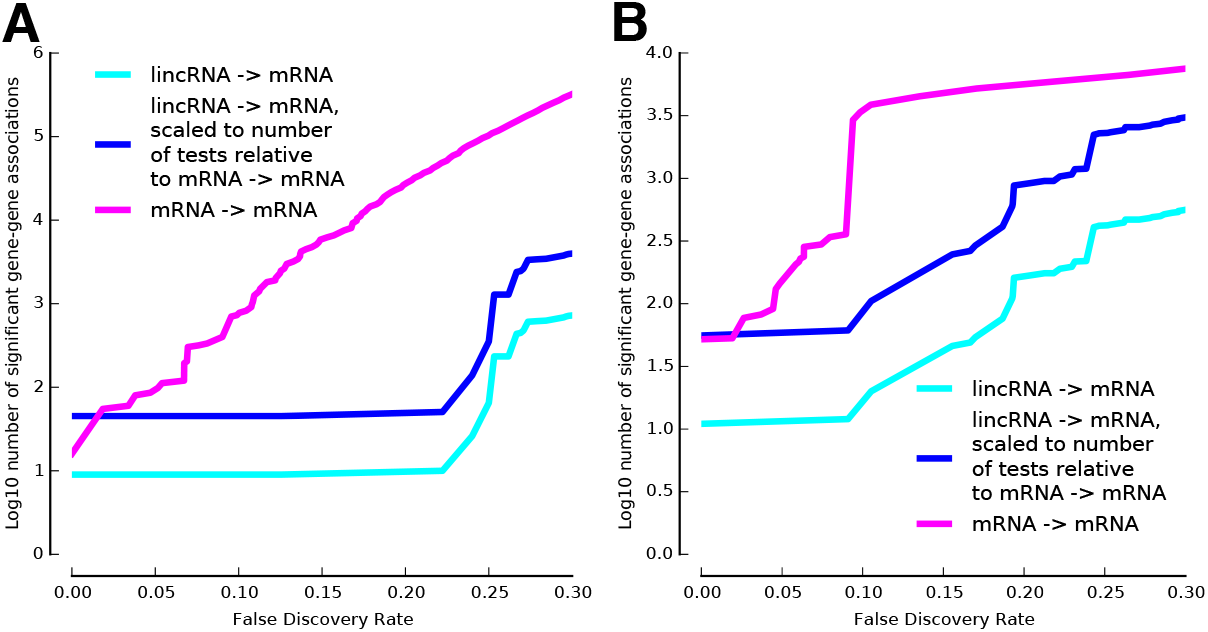
Number of discoveries of trans-associations by false discovery rate. **Panel A**, Number of trans gene-gene associations discovered by Mendelian randomization by FDR threshold. **Panel B**, Number of trans eQTL-gene associations discovered by using the best cis-eQTLs per gene as candidate trans eQTLs by FDR threshold.

#### 2.6.2 Mendelian randomization approach for testing for trans-effects of lincRNAs

To study whether lincRNAs affect mRNA expression levels genome-wide in comparison with mRNA in the presence of shared confounding effects, we developed and applied a Mendelian randomization (MR) test to quantify support a direct relationship between each RNA that has a cis-eQTL and all local and distal genes.

Mendelian randomization (MR) is a form of causal inference referred to as instrumental variable analysis, where an *instrumental variable* (here, a cis-eQTL) is used to artificially create randomization from observational data, leading to an analysis that mimics a controlled randomized trial (CRT) [94]. CRT analyses are the gold standard approach to testing for causal effects, so MR is a powerful approach to causal inference in observational genomic data. MR requires that a genotype (instrument) is associated with the independent variable (the cis-regulated gene; Fig. 6). MR explicitly controls for unobserved co-regulators of the two genes via the instrumental variable. In particular, our approach to MR separates variation in the independent variable *x* (the cis-gene) due to the cis-eQTL and variation due to all other effects, and only uses the genetically-regulated variation to test for association with the dependent variable (the trans-gene).

The effect sizes of 1,077 best cis-linc-eQTLs and 5,899 best cis-pc-eQTLs were compared to the effect sizes of those same eQTLs in trans by means of Mendelian randomization. We found that 36 (3.3% of tested) lincRNA genes associated with the expression of 753 protein-coding genes, while 278 (4.7% of tested) protein-coding genes associated with the expression of 4, 298 protein-coding genes (20% FDR; FET, *p* = 4.57 × 10^−2^; Fig. 7B). While the proportion of lincRNA genes with trans associations was significantly lesser than the proportion of protein-coding genes with trans associations, our results support the hypothesis that lincRNAs participate in a dosage-dependent distal regulation of protein coding gene transcription on a genome-wide scale.

We suspect that it is the additional control in MR for shared confounding effects that leads to different conclusions between the näive approach and the MR approach. This effect will be compounded in the shared batch and technical effects in the measurements of the cis- and trans-genes expression levels, as the assays were shared. We also note that the tests were implemented differently: the MR approach combined gene expression levels across all tissues, whereas the näive approach explicitly used cross-tissue meta-analyses.

To ensure that trans-pc-eQTLs were not long-range cis-pc-eQTLs, we verified that trans-pc-eQTLs from cis-pc-eQTLs were located on the same chromosome as their associated genes at a frequency no greater than expected by chance, when modeling the probability of choosing a SNP or gene as a uniform distribution across the length of each chromosome (Z-test, *p* > 0.5).

Previous work has speculated that lincRNAs, and lncRNAs more generally, compose a novel layer of transcription regulation [37, 63, 88]. Researchers have experimentally shown that a handful of lincRNAs have regulatory functions, for example, in *Mus musculus* embryonic stem (ES) cell differentiation [37] and in human lung endoderm morphogenesis [42]. Certain lincRNAs with established regulatory functions did not have cis-linc-eQTLs in our data set and thus could not be tested for trans-pc-eQTL association. These excluded functional lincRNAs include *TERC, Xist, lincRNA-ROR, MEG3, MEG8, ncRNA-a1, ncRNA-a5*, and *ncRNA-a7*. Another two lincRNAs, *NEAT1* and *MALAT1/NEAT2*, which have been hypothesized to have trans regulatory functions in humans based on experimental characterization of their cellular function, did have cis-linc-eQTLs in our study (166 and 50, respectively). However, *NEAT1* and *MALAT1/NEAT2* did not significantly associate with the expression of any protein-coding genes in trans. Many of the other well-studied lncRNAs with known regulatory roles are either classified as antisense RNA, miscRNA, or were not annotated in GENCODE and so were not tested here. These lncRNAs include *HOTAIR, HOTTIP, Evf2, Kcnq1ot1, Paupar, CCND1*-associated ncRNAs, *AIR, DHFR* minor promoter, *NRON, ANRIL, ncRNA-a2, ncRNA-a3, ncRNA-a4*, and others.

### 2.7 Local correlation of mRNAs and lincRNAs

A handful of lincRNAs has been shown to regulate mRNA expression both locally and distally by diverse mechanisms [39]. Mendelian randomization assumes that the cis-eQTL does not have a direct effect on the downstream gene, which we could not guarantee in this scenario, but we suspect that direct distal effects will be uncommon.

To test for local genomic interaction between lincRNAs and mRNAs, we used correlation coefficients to test for co-regulation of gene pairs. The median absolute correlation coefficient (MACC) was significantly higher for gene pairs that were nearer to one another compared to those farther away (Mann-Whitney U-test, 20 Kb versus 100 Kb: *p* ≤ 1 × 10^−100^; 100 Kb versus 1 Mb: *p* ≤ 1 × 10^−100^; 1 Mb versus 5 Mb: *p* ≤ 1 × 10^−100^). However, across all gene pair genomic proximity thresholds, the MACC between pairs of lincRNAs and protein-coding genes was not significantly greater than the MACC for pairs of protein-coding genes nor for pairs of lincRNA genes (Mann-Whitney U-test, *p* > 0.05). In other words, proximal pairs of lincRNAs and protein-coding genes showed no more correlation in expression than other proximal gene pairs, as has been shown in previous work [15, 99].

## 3 Discussion

In this study, we applied a Mendelian randomization approach to explore the biological role of lincRNAs through their associated genetic variants, using matched-power analyses on mRNAs as an experimental control. We provided evidence for similar allele-specific regulation of gene expression levels among lincRNA and mRNA genes using CRE enrichment analyses. We find that organismal traits—here, largely adipose-related traits—may be genetically mediated through lincRNA expression levels; we compiled 74 explanatory lincRNAs and highlight six of the most promising lincRNAs that may be involved in the regulation of specific organismal traits. Using Mendelian randomization, we found that there is evidence for lincRNAs having distal effects on the expression levels of protein-coding genes in a dosage-dependent way across the genome in similar proportions to distal effects of mRNA.

Mendelian randomization shows great promise in performing genome-wide studies of a functional role for specific types of epigenetic elements. We contrasted the results using MR with results using standard tests of trans-eQTLs, where the discovery set of SNPs included only cis-eQTLs. The improvement in results of MR was clear: we found diminishingly few trans-eQTL associations from cis-pc-eQTLs (39) and cis-linc-eQTLs (4) in the univariate analysis tissue-by-tissue. Using MR and pooling data across tissues, however, we found 5, 213 trans gene-gene associations (from 278 putative regulators; FDR ≤ 20). There are a number of limitations to our study, including a small sample size for eQTL discovery, noisy lincRNA annotations in the human genome as compared with annotations of mRNA, and lack of experimental approaches to validating trans-eQTLs that act via lincRNAs. None of our results, however, are conditioned on these limitations.

We designed our study to have similar statistical power to detect both the effects of lincRNA regulation of mRNA and mRNA regulation of mRNA. This controlled experiment was possible since we used a matched distribution of effect sizes of the cis-eQTLs and the use of a MR approach to quantify the magnitude of the effects of the cis-RNA on all trans-mRNA to control for shared confounding effects. We proposed a specific MR test to evaluate distal regulatory effects in the context of this controlled experiment. We feel that these methods can be generalized to increase statistical power in trans-eQTL studies [104], study genetic mechanisms of specific complex traits [33], or infer the causal relationships among epigenetic markers [8]. Here we apply this approach to assess, at a genome-wide scale, the regulatory potential of non-coding RNA - specifically to determine whether or not lincRNAs have distal regulatory effects.

Methodologically, there are many avenues to pursue in the area of methods design for Mendelian randomization. In particular, we are scaling our approach to high-dimensional SNPs and epigenetic markers. Our current method, implemented in an open source Python package on GitHub (https://github.com/PrincetonUniversity/MReQTL), can be applied to data for which summary statistics are available across unpaired samples; in other words, we did not leverage the paired samples here, and we could have equivalently ascertained lincRNA cis-eQTLs in one dataset and mRNA trans-eQTLs in a second dataset. A paired test that simultaneously controls for unobserved confounding and exploits the paired sample design to improve statistical power would be valuable in light of the many paired epigenetic studies on the horizon.

## 4 Methods

### 4.1 RNA-seq processing, mapping, quantification, and normalization

RNA-seq reads from the Genotype-Tissue Expression (GTEx) Project consortium Pilot data [58] were trimmed using Trimmomatic (v.0.30) [11]. We mapped reads to human reference genome assembly GRCh37.p13 using STAR aligner (v.2.3.0) [26]. Of 15.93 B total read pairs mapped, 14.52 B read pairs (91%) mapped uniquely, and 1.2 B read pairs (7.5%) were discarded for mapping to more than one locus. Mapped reads were converted into read counts (one read pair per read count) using the software featureCounts [55] on settings that required both ends of a read pair to be mapped to the same gene. For all further analyses, we used expression levels for lincRNAs and mRNAs in autosomal chromosomes with non-zero expression levels in at least 10% of samples in at least one tissue, which filtered 32% of the original 6, 416 lincRNA genes and 10.6% of the original 20, 345 protein-coding genes. After filtering, we retained 4,377 lincRNA genes and 18, 194 protein-coding genes. Read abundances for lincRNA and protein-coding genes were normalized by gene length, GC-content, and library size using the Bioconductor R package cqn [40], which yielded RPKM estimates. We controlled for known and unknown covariates of expression by removing all principal components that explained greater than 0.5% of the variation in expression values across individuals. Principal components analysis was performed using NumPy and SciPy Python packages. Expression values were normalized to the quantiles of the standard normal distribution across individuals within gene and tissue before and after the removal of PCs.

### 4.2 RNA-seq read trimming and preprocessing

Adapter sequences and overrepresented contaminant sequences, identified by FastQC (v.0.10.1) [4], were trimmed using Trimmomatic (v.0.30) [11] with 2 seed mismatches and a simple clip threshold of 20. Leading and trailing nucleotides (low quality or Ns) were trimmed from all reads until a canonical base was encountered with quality greater than 3. For adaptive quality trimming, reads were scanned with a 4-base sliding window, trimming when the average quality per base dropped below 20. Any remaining sequences shorter than 30 nucleotides were discarded. After trimming, the original raw read counts of 5.87 B read pairs in adipose, 6.34 B read pairs in artery, 7.89 B read pairs in lung, 6.45 B read pairs in skin were reduced to 3.65 B read pairs in adipose, 4.02 B read pairs in artery, 4.63 B read pairs in lung, 3.63 B read pairs in skin.

### 4.3 RNA-seq read mapping

After preparing the genome with STAR aligner genomeGenerate mode using a splice junction database (sjdbGTFfile) set to GENCODE v.19 annotation, the splice junction database overhang (sjdbOverhang) set to 75 bp, and defaults for all remaining settings. STAR aligner alignReads mode was run using default settings except outFilterMultimapNmax was set to 1 so that only uniquely mapping reads were retained.

### 4.4 Filtering and imputing genotype data

In the same subjects from which RNA-seq samples were taken, 3.5 M quality-filtered SNPs were assayed on the Illumina Omni5-Quad array. These SNPs were then used to impute to approximately 10 M SNPs using as reference the 1000 Genomes phase 1 release [1] as described in earlier work [58]. Imputed SNPs with quality score < 0.4 and diverging from Hardy-Weinberg equilibrium (*p* ≤ 1 × 10^−6^) were removed before association mapping.

### 4.5 Association mapping for cis-effects

Bayesian regression was performed using two separate techniques, eQTLBMA (v. 1.3) [30] and SNPTEST (v. 2.5.b.4). eQTLBMA was used to perform regression jointly on data from all four tissue types, which enabled the estimation of the proportion of expression quantitative trait loci (*eQTLs*) shared across tissues and allowed for greater power than would be possible by performing regression for each tissue separately.

The parameters we used corresponded to the *BF_BMA_* as previously described [30]. After sample test runs, we filtered SNPs with minor allele frequency (MAF) < 0.02 as low MAF SNPs were found to be uninformative when using eQTLBMA. Using SNPTEST, we assumed an additive effects model with a prior effect size modeled as a normal distribution with zero mean and variance modeled as an inverse gamma scaled by a factor of 0.02 and with mean 3 and variance 2. We tested associations between the expression of each lincRNA or mRNA and all SNPs in the interval between 100 Kb upstream of the transcription start site (TSS) to 100 Kb downstream of the transcription end site (TES). For comparison, *local* association mapping was also performed for a candidate region extending from 1 Mb upstream and downstream of each TSS and TES respectively. For each tested gene-SNP pair, eQTLBMA returned Laplace approximations of log_10_ Bayes factors (log_10_BFs) [30] and SNPTEST also returned approximate log_10_BFs for the observed expression-genotype relationships and for expression-genotype relationships where labels of individuals were randomly permuted. Because of the computational burden, only one permutation was performed for both SNPTEST and eQTLBMA. (For numbers of genes, SNPs and gene-SNPs tested for association, see Supplemental Table 1). The false discovery rate (FDR) was computed as the number of associations identified in the permuted data at each log_10_BF cutoff value divided by the number of associations identified in the observed data at that same cutoff, as in previous work [60]. Significant cis-eQTLs—both lincRNA (*linc-eQTLs*) and protein-coding (*pc-eQTLs*)—were defined as those gene-SNP associations that passed the significance threshold at a 5% FDR.

### 4.6 Distribution of location of the best cis-eQTLs

For all lincRNA and protein-coding genes with at least one cis-eQTL, the upstream and downstream cis association candidate regions flanking the gene body were divided into ten equal parts into which the best eQTLs were binned (e.g. if eQTL was 100, 000 - 90, 000 bp upstream, then bin 1, if eQTL was 89, 999 - 80, 000 bp upstream, then bin 2, etc.) Gene bodies were split into different numbers of bins according to the mean length of genes in the class under consideration such that the genic bins would be roughly 10 Kb in size, and thus comparable to the flanking bins. This worked out to 9 bins per gene for SNPTEST pc-eQTLs, 7 bins per gene for eQTLBMA pc-eQTLs, 4 bins per gene for SNPTEST linc-eQTLs, and 3 bins per gene for eQTLBMA linc-eQTLs.

### 4.7 Comparison of cis-linc-eQTLs and cis-pc-eQTLs in tissue specificity

To determine whether linc-eQTLs were more tissue-specific than pc-eQTLs, we modeled the probability that an eQTL was tissue-specific (i.e., significant in only one of the four tissues) as a logistic regression model with the following covariates: eQTL type (linc-eQTL or pc-eQTL), absolute effect size, log of distance to TSS/TES, log of average expression level, MAF, log of the transcribed gene length, log of the genomic gene length, tissue specificity of expression score, GC-content, and log of the number of exons.

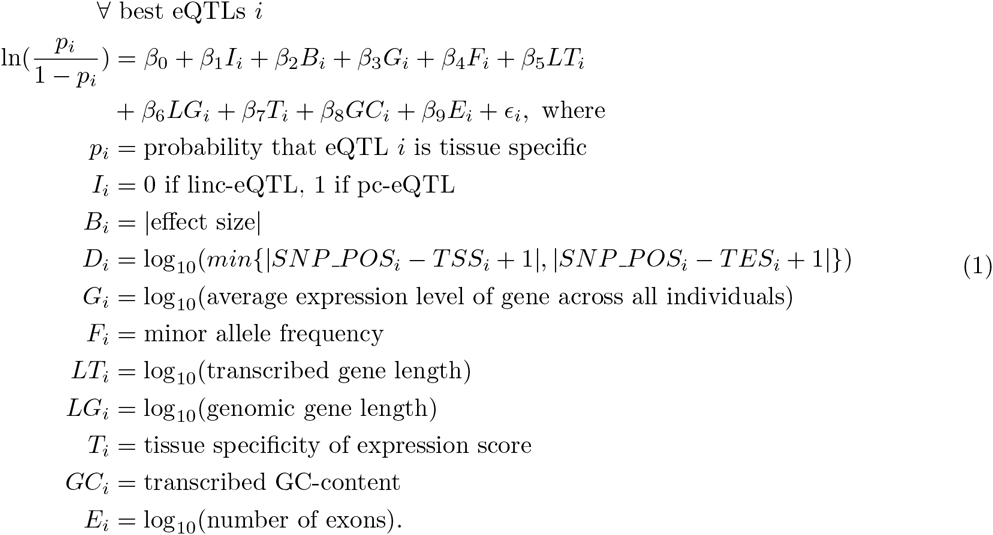

### 4.8 Conditional analysis of shared cis-linc-/pc-eQTLs

We allowed for four different scenarios of the regulatory relationship between cis-eQTL, lincRNA, and mRNA for those cis-eQTLs that were both linc-eQTLs and pc-eQTLs:

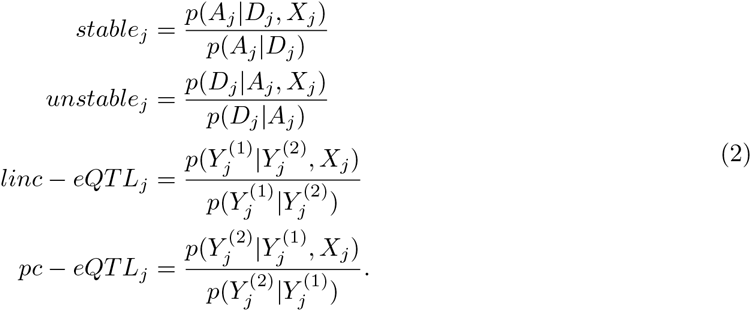

In each case, we modeled the probability using a Bayesian linear model:

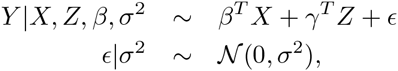

where the prior on effect size is the D2 prior from previous work [92]. We summarized this conditional analysis with a modified Bayes factor (BF) that is computed as follows for SNP *j*:

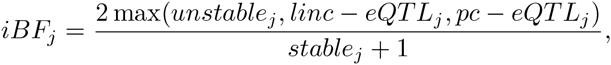

where 1 in the denominator captures the null (*none_j_*) hypothesis in the denominator of all of these separate tests. When *iBF_j_* > 1, there is evidence that the eQTL effect is differential; otherwise it is stable or not present.

### 4.9 Enrichment for overlap with cis-regulatory elements

Significant cis-linc-eQTLs and cis-pc-eQTLs were tested for enrichment of overlap with various cis-regulatory elements (CREs) in matching ENCODE [20] cell lines. The background distribution consisted of all SNPs tested for association that had MAFs at least as great as the smallest MAF for any significant linc-eQTL or pc-eQTL and nearest to a GENCODE v.19 protein-coding or lincRNA gene with non-zero average expression. There were 2.0 M of these best linc-eQTL background SNPs, 3.0 M for all linc-eQTL background SNPs, 5.8 M for best pc-eQTL background SNPs, and 7.6 M for all pc-eQTL background SNPs. The probability of a SNP overlapping a CRE was modeled in a logistic regression model, including as covariates the logarithm of the distance of SNP to associated gene, average expression levels of the associated gene, MAF, and an indicator of whether the SNP was an cis-eQTL (Eqn. 4). As background SNPs could not be assigned an associated gene (by definition, such SNPs have no associations), they were assigned to the nearest gene, breaking ties by choosing the nearest gene with the highest average expression.

The calculation of bootstrap confidence intervals for odds ratio of probability of overlap with CRE and odds ratio of probability of linkage to trait-associated SNP throughout this work refers to the bias-corrected accelerated bootstrap method [27] and was implemented using `scikit.bootstrap` [76].

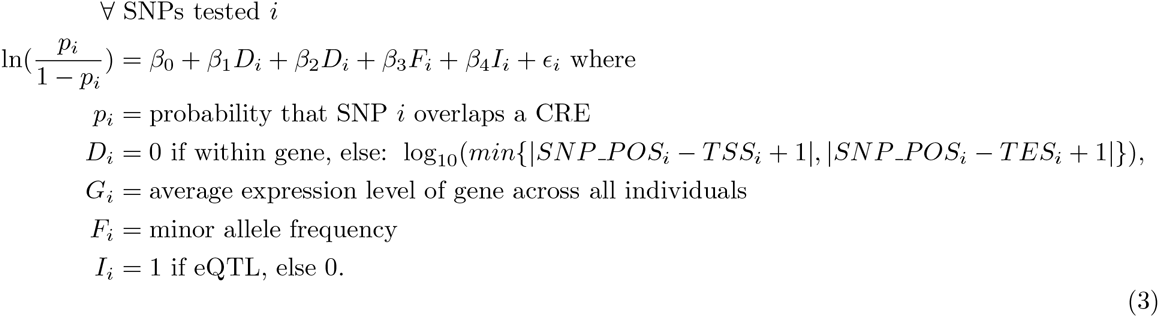

### 4.10 Enrichment for linkage to trait-associated SNPs (TASs)

Cis-linc-eQTLs and cis-pc-eQTLs were tested for enrichment of linkage to TASs. Genome-wide assocication study (GWAS) SNPs were downloaded from the NHGRI GWAS catalog (accessed: 04/22/2014) [103]. The background set of SNPs was compiled as described above in enrichment for overlap with cis-regulatory elements. Linkage disequilibrium (LD), or the non-random association of alleles due to shared ancestry, can be measured by the correlation coefficient (*r*^2^) between loci. A SNP was considered linked to a TAS if the SNP and the TAS were located within 100 Kb and had a pairwise *r*^2^ ≥ 0.8. The probability of a SNP being linked to a TAS was modeled in a logistic regression (shown below) as a function of the logarithm of distance of SNP to associated gene, average expression of the associated gene, MAF, and an indicator of whether the SNP was a cis-eQTL. As background SNPs could not be assigned an associated gene (by definition, such SNPs have no associations), they were assigned to the nearest gene, breaking ties by choosing the nearest gene with the highest average expression (Eqn. 5).

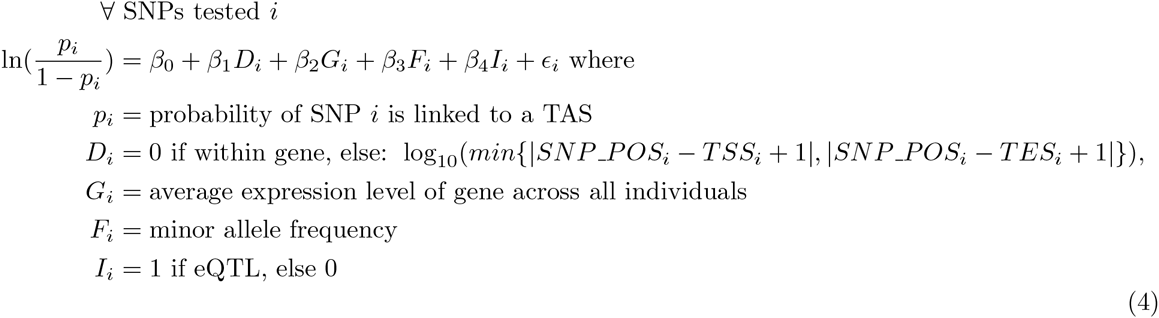

### 4.11 Evaluating mechanistic hypotheses for explanatory lincRNAs

We compiled a list of TASs that may regulate the associated trait through the regulation of lincRNA expression levels. For this set of *lincRNA-TASs*, we selected TASs that were at least 10 Kb away from the nearest GENCODE v.19 protein-coding gene and were in LD with a cis-linc-eQTLs (*r*^2^ > 0.8) but were not in LD with a cis-pc-eQTLs or a local-pc-eQTL (*r*^2^ < 0.2) (Supplemental Table 4). The list of explanatory lincRNAs were those lincRNAs associated with lincRNA-TASs as described above (Supplemental Table 3). To study the enrichment of certain trait types in the set of lincRNA-TASs, we compared the distribution of traits for lincRNA-TASs to the distribution of traits for TASs linked to pc-eQTLs and not to linc-eQTLs. To study a wide range of possible mechanisms of action for the lincRNAs on an organismal phenotype, we tested the candidate set of lincRNAs for genetic signatures of potential mechanisms of action.

To study a wide range of possible mechanisms of action for the lincRNAs on an organismal phenotype, we tested the candidate set of lincRNAs for genetic signatures of potential mechanisms of action. In particular, we tested:

- *coding potential* as estimated by CPAT (v1.2.1) [101],
- *sequence conservation* as estimated by phastCons (46-way) and downloaded from the UCSC Genome Browser (accessed: 12/18/2014) [93],
- *conserved secondary structure* as estimated by Evofold and downloaded from the UCSC Genome Browser (accessed: 12/19/2014) [75],
- *secondary structure folding energy* as estimated by RNAfold (v.2.1.3) [59],
- *number of miRNA binding sites* as estimated by TargetScan [54] and compiled in lnCeDB (accessed: 01/14/2015) [21],
- *miRNA binding site conservation* (46-way and primates-only) as estimated by TargetScan [54] and compiled in miRcode database (accessed: 01/14/2015) [48],
- *nuclear versus cytoplasm localization ratio* as measured for selected GENCODE v.7 transcripts [23],
- *signal peptide potential* as measured by signalP (v.4.1) [77].

For the coding potential analysis, the sequences of all lincRNAs tested for linc-eQTLs were compared to their reverse complement RNA sequences, which are assumed to have coding potential equivalent to genomic background but have identical length distribution as the set of lincRNAs, representing our null hypothesis. For other analyses, metrics for candidate lincRNAs were compared to their complement set among all lincRNAs tested. For those metrics that were continuous or integer-valued—coding potential, secondary structure folding energy, number of miRNA binding sites, conservation of miRNA binding sites, signal peptide potential, nuclear versus cytoplasm localization ratio—the metrics were compared between sets using a Mann-Whitney U-test. For conserved secondary structure, which was a binary metric, the metrics were compared between sets using Fisher’s exact test. Those metrics that could be reasonably binarized—coding potential, conservation of miRNA binding sites with varied thresholds, signal peptide potential, nuclear versus cytoplasm localization ratio ≥ 1 versus < 1—the metrics were binarized within set and were compared across sets using Fisher’s exact test. Any test that yielded significant results using the methods above was retested using multivariate linear regression or logistic regression to control for transcript length, which influences many of the above metrics.

### 4.12 Allele-specific luciferase assay

We selected for validation cis-eQTLs that were located in DHS shared across cell types that represented the tissues used: the lung epithelial cell line A549, primary preadipocytes, the skin fibroblast cell line AG04449, and aortic adventitial fibroblasts [20]. We used PCR to amplify the major and minor haplotype of each DHS from the genomes of 1,000 Genomes Project donors [1]. (Details of the target regions, haplotypes, and primers used are in Supplemental Table 5.) The DHS were ligated upstream of the Simian virus 40 promoter driving expression of a firefly-derived luciferase reporter gene (Promega pGL4.13) using Gibson assembly. The constructs were then transformed into *Escherichia coli* (*E. coli*) and haplotypes were confirmed with Sanger sequencing. The positive clones were expanded, and plasmids purified using standard approaches. We then transiently transfected each luciferase reporter into HepG2 cells, and calculated the relative luciferase expression between the two haplotypes after normalizing to a renilla-derived luciferase expressed from a co-transfected control plasmid. The statistical significance of differences in renilla-normalized enhancer activity between haplotypes was evaluated using a Student’s t-test.

### 4.13 Association mapping for trans-effects

We performed association mapping (both single-tissue and multi-tissue) for all lincRNA and protein-coding gene expression levels (tested in cis) in all tissue types using only the 2, 034 best cis-linc-eQTLs and 13, 522 best cis-pc-eQTLs, which had the effect of greatly reducing the multiple hypothesis testing burden to detect trans-eQTLs. To exclude any cis-associations, we did not test for trans effects for any SNPs within 1 Mb of a gene. The same parameters were used for SNPTEST and eQTLBMA trans-association as described above for cis-association mapping. As in cis-association eQTL mapping, one permutation was performed for trans-eQTL analyses. A significance threshold was determined empirically corresponding to a 20% FDR.

### 4.14 Mendelian randomization

We used an instrumental variable (IV) analysis to identify cis-regulated genes (both mRNA and lincRNA) that affect regulation of protein coding genes in trans. Let *z* be the *instrumental variable* (here, the genotype of a SNP across *n* indivduals) that directly affects the *independent variable x* (here, the cis-RNA), which, in turn, may or may not affect the *dependent variable y* (here, the trans-mRNA). Both *x* and *y* may be affected by unmeasured variables *u* (Fig. 6). Mendelian randomization (MR) is an example of IV analysis, where we quantify the degree of the causal relationship from *x* to *y* by removing all sources of variation in *x* other than variation due to instrumental variable *z* and then testing for a relationship between this *denoised x* and *y*. The MR framework requires *z* to have direct effects on *x* and indirect effects only on *y*, and also implicitly controls for all possible unmeasured confounders *u* that jointly affect *x* and *y*.

First, define the normal equation for linear regression, which estimates the effect size of a linear regression predictor *x* on a response variable *y:* 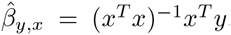. The linear regression model, 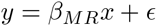, may be conditioned on the instrumental variable *z*, and then expectations taken:

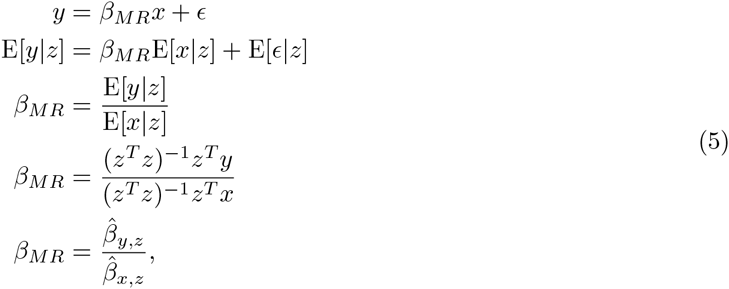

where 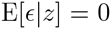 by assumption and 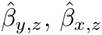 are the estimated effect sizes for the trans-eQTL and the cis-eQTL respectively. Thus, to quantify the direct effects of RNA *x* on mRNA *y* using *β_MR_*, we compute the ratio of the trans-eQTL to the cis-eQTL.

We now test the null hypothesis, that *β_MR_* = 0, compared to the alternative hypothesis, 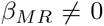, using Wald’s test. The test statistic, 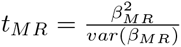 has a χ^2^ distribution with three degrees of freedom (two linear regression intercepts and two slopes). We compute the variance terms, *var*(*β_MR_*) and residual variance *σ*^2^, as follows:

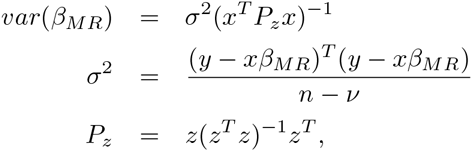

where *ν* is the degrees of freedom and *n* is the number of samples.

We assessed global FDR using a permutation. In particular, the null hypothesis is that there is no relationship between *x* and *y*. Correspondingly, we used the estimated effect size 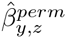 from permuted dependent variable data while maintaining the estimated effect sizes 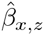 from the unpermuted samples to ensure there is still an association between *z* and *x*. Then, using the ordered list of p-values computed using Wald’s test, we approximated the global FDR for specific p-value thresholds using the number of p-values in the permutations that were smaller than the threshold (FPs) divided by the number of p-values in the true data that were smaller than the threshold (TPs+FPs).

### 4.15 Local correlation of mRNA and lincRNA

We computed pairwise Pearson’s correlation coefficients between gene expression estimates across all samples from the four tissue types for gene pairs within multiple genomic proximity thresholds (20 Kb, 100 Kb, 1 Mb, 5 Mb) using the NumPy function corrcoef.

## Acknowledgements

BEE was funded by NIH R00 HG006265 and NIH R01 MH101822. CDB was funded by NIH R01 MH101822. AAP was supported by a postdoctoral fellowship from the Jane Coffin Childs Foundation. CG, CMV, and TER were funded by NIH R01 DK099820. The data reported in this paper are tabulated in the Supporting Online Material and archived at dbGaP, study accession number phs000424.v4.p1, Common Fund Genotype-Tissue Expression (GTEx) Project.

## Author Contributions

B.E.E., T.E.R., and I.C.M. conceived the experiments. B.E.E., and I.C.M. designed the experiments. I.C.M. performed the computational experiments. I.C.M. and B.E.E. analyzed the data. C.G., C.M.V., and T.E.R. designed and performed the laboratory experiments. I.C.M., A.A.P., C.D.B., T.E.R., and B.E.E. wrote the paper.

## Disclosure Declaration

The authors have no conflicts of interests to disclose.

## References

1. 1000 Genomes Project Consortium, 2010. A map of human genome variation from population-scale sequencing. Nature, 467(7319):1061–1073.

2. Anderson D. M., Anderson K. M., Chang C.-L., Makarewich C. A., Nelson B. R., McAnally J. R., Kasaragod P., Shelton J. M., Liou J., Bassel-Duby R., et al., 2015a. A micropeptide encoded by a putative long noncoding rna regulates muscle performance. Cell, 160(4):595–606.

3. Anderson D. M., Anderson K. M., Chang C.-L., Makarewich C. A., Nelson B. R., McAnally J. R., Kasaragod P., Shelton J. M., Liou J., Bassel-Duby R., et al., 2015b. A micropeptide encoded by a putative long noncoding rna regulates muscle performance. Cell, 160(4):595–606.

4. Andrews S., 2012. http://www.bioinformatics.bbsrc.ac.uk/projects/fastqc/.

5. arcOGEN Consortium et al., 2012. Identification of new susceptibility loci for osteoarthritis (arcOGEN): a genome-wide association study. The Lancet, 380(9844):815–823.

6. Ardlie K. G., Deluca D. S., Segre A. V., Sullivan T. J., Young T. R., Gelfand E. T., Trowbridge C. A., Maller J. B., Tukiainen T., Lek M., et al., 2015. The Genotype-Tissue Expression (GTEx) pilot analysis: Multitissue gene regulation in humans. Science, 348(6235):648–660.

7. Azzalin C. M., Reichenbach P., Khoriauli L., Giulotto E., and Lingner J., 2007. Telomeric repeat–containing rna and rna surveillance factors at mammalian chromosome ends. Science, 318(5851):798–801.

8. Banovich N., Lan X., McVicker G., van de Geijn, B., Degner J., Blischak J., Roux J., Pritchard J., and Gilad Y., 2014. Methylation QTLs are associated with coordinated changes in transcription factor binding, histone modifications, and gene expression levels. PLoS Genetics, 10(9):e1004663–e1004663.

9. Bassett A. R., Akhtar A., Barlow D. P., Bird A. P., Brockdorff N., Duboule D., Ephrussi A., Ferguson-Smith, A.C., Gingeras T. R., Haerty W., et al., 2014. Considerations when investigating lncRNA function in vivo. eLife, 3:e03058.

10. Bazzini A. A., Johnstone T. G., Christiano R., Mackowiak S. D., Obermayer B., Fleming E. S., Vejnar C. E., Lee M. T., Rajewsky N., Walther T. C., et al., 2014. Identification of small ORFs in vertebrates using ribosome footprinting and evolutionary conservation. The EMBO Journal, 33(9):981–993.

11. Bolger A. M., Lohse M., and Usadel B., 2014. Trimmomatic: a flexible trimmer for Illumina sequence data. Bioinformatics.

12. Brown C. D., Mangravite L. M., and Engelhardt B. E., 2013. Integrative Modeling of eQTLs and Cis-Regulatory Elements Suggests Mechanisms Underlying Cell Type Specificity of eQTLs. PLoS Genetics, 9(8):e1003649.

13. Brown C. J., Hendrich B. D., Rupert J. L., Lafreniere R. G., Xing Y., Lawrence J., and Willard H. F., 1992. The human *XIST* gene: Analysis of a 17 kb inactive X-specific RNA that contains conserved repeats and is highly localized within the nucleus. Cell, 71(3):527–542.

14. Buiting K., Nazlican H., Galetzka D., Wawrzik M., Groβ S., and Horsthemke B., 2007. C15orf2 and a novel noncoding transcript from the prader–willi/angelman syndrome region show monoallelic expression in fetal brain. Genomics, 89(5):588–595.

15. Cabili M. N., Trapnell C., Goff L., Koziol M., Tazon-Vega B., Regev A., and Rinn J. L., 2011. Integrative annotation of human large intergenic noncoding RNAs reveals global properties and specific subclasses. Genes Dev., 25(18):1915–1927.

16. Choi J., Southworth L. K., Sarin K. Y., Venteicher A. S., Ma W., Chang W., Cheung P., Jun S., Artandi M. K., Shah N., et al., 2008. Tert promotes epithelial proliferation through transcriptional control of a myc-and wnt-related developmental program. PLoS Genetics, 4(1):e10.

17. Chu C., Qu K., Zhong F. L., Artandi S. E., and Chang H. Y., 2011. Genomic maps of long noncoding rna occupancy reveal principles of rna-chromatin interactions. Molecular cell, 44(4):667–678.

18. Clemson C. M., McNeil J. A., Willard H. F., and Lawrence J. B., 1996. *XIST* RNA paints the inactive × chromosome at interphase: evidence for a novel RNA involved in nuclear/chromosome structure. The Journal of Cell Biology, 132(3):259–275.

19. Comuzzie A. G., Cole S. A., Laston S. L., Voruganti V. S., Haack K., Gibbs R. A., and Butte N. F., 2012. Novel genetic loci identified for the pathophysiology of childhood obesity in the hispanic population. PloS One, 7(12):e51954.

20. Consortium E. P. et al., 2012. An integrated encyclopedia of DNA elements in the human genome. Nature, 489(7414):57–74.

21. Das S., Ghosal S., Sen R., and Chakrabarti J., 2014. lnCeDB: database of human long noncoding RNA acting as competing endogenous RNA. PloS one, 9(6):e98965.

22. Dastani Z., Hivert M.-F., Timpson N., Perry J. R., Yuan X., Scott R. A., Henneman P., Heid I. M., Kizer J. R., Lyytikäinen L.-P., et al., 2012. Novel loci for adiponectin levels and their influence on type 2 diabetes and metabolic traits: a multi-ethnic meta-analysis of 45,891 individuals. PLoS Genetics, 8(3):e1002607.

23. Derrien T., Johnson R., Bussotti G., Tanzer A., Djebali S., Tilgner H., Guernec G., Martin D., Merkel A., Knowles D. G., et al., 2012. The GENCODE v7 catalog of human long noncoding RNAs: Analysis of their gene structure, evolution, and expression. Genome Research, 22(9):1775–1789.

24. Dimas A. S., Deutsch S., Stranger B. E., Montgomery S. B., Borel C., Attar-Cohen H., Ingle C., Beazley C., Arcelus M. G., Sekowska M., et al., 2009. Common regulatory variation impacts gene expression in a cell type–dependent manner. Science, 325(5945):1246–1250.

25. Dimitrova N., Zamudio J. R., Jong R. M., Soukup D., Resnick R., Sarma K., Ward A. J., Raj A., Lee J. T., Sharp P. A., et al., 2014. *LincRNA-p21* activates p21 in cis to promote Polycomb target gene expression and to enforce the G1/S checkpoint. Molecular Cell, 54(5):777–790.

26. Dobin A., Davis C. A., Schlesinger F., Drenkow J., Zaleski C., Jha S., Batut P., Chaisson M., and Gingeras T. R., 2013. STAR: ultrafast universal RNA-seq aligner. Bioinformatics, 29(1):15–21.

27. Efron B., 1987. Better bootstrap confidence intervals. The Journal of the American Statistical Association, 82(397):171–185.

28. Emilsson V., Thorleifsson G., Zhang B., Leonardson A. S., Zink F., Zhu J., Carlson S., Helgason A., Walters G. B., Gunnarsdottir S., et al., 2008. Genetics of gene expression and its effect on disease. Nature, 452(7186):423–428.

29. Fairfax B. P., Makino S., Radhakrishnan J., Plant K., Leslie S., Dilthey A., Ellis P., Langford C., Vannberg F. O., and Knight J. C., et al., 2012. Genetics of gene expression in primary immune cells identifies cell type-specific master regulators and roles of hla alleles. Nature Genetics, 44(5):502–510.

30. Flutre T., Wen X., Pritchard J., and Stephens M., 2013. A statistical framework for joint eQTL analysis in multiple tissues. PLoS Genetics, 9(5):e1003486.

31. Fox C. S., Liu Y., White C. C., Feitosa M., Smith A. V., Heard-Costa N., Lohman K., Johnson A. D., Foster M. C., Greenawalt D. M., et al., 2012. Genome-wide association for abdominal subcutaneous and visceral adipose reveals a novel locus for visceral fat in women. PLoS Genetics, 8(5):e1002695.

32. Gaffney D. J., Veyrieras J.-B., Degner J. F., Pique-Regi R., Pai A. A., Crawford G. E., Stephens M., Gilad Y., and Pritchard J. K., 2012. Dissecting the regulatory architecture of gene expression QTLs. Genome Biology, 13(1):R7.

33. Gamazon E. R., Wheeler H. E., Shah K. P., Mozaffari S. V., Aquino-Michaels K., Carroll R. J., Eyler A. E., Denny J. C., Nicolae D. L., Cox N. J., et al., 2015. A gene-based association method for mapping traits using reference transcriptome data. Nature Genetics, 47(9):1091–1098.

34. Grote P., Wittler L., Hendrix D., Koch F., Währisch S., Beisaw A., Macura K., Bläss G., Kellis M., Werber M., et al., 2013. The tissue-specific lncRNA *Fendrr* is an essential regulator of heart and body wall development in the mouse. Developmental Cell, 24(2):206–214.

35. Grundberg E., Small K. S., Hedman, Å. K., Nica A. C., Buil A., Keildson S., Bell J. T., Yang T.-P., Meduri E., Barrett A., et al., 2012. Mapping cis-and trans-regulatory effects across multiple tissues in twins. Nature Genetics, 44(10):1084–1089.

36. Guttman M., Amit I., Garber M., French C., Lin M. F., Feldser D., Huarte M., Zuk O., Carey B. W., Cassady J. P., et al., 2009. Chromatin signature reveals over a thousand highly conserved large non-coding RNAs in mammals. Nature, 458(7235):223–7.

37. Guttman M., Donaghey J., Carey B. W., Garber M., Grenier J. K., Munson G., Young G., Lucas A. B., Ach R., Bruhn L., et al., 2011. lincrnas act in the circuitry controlling pluripotency and differentiation. Nature, 477(7364):295–300.

38. Guttman M., Garber M., Levin J. Z., Donaghey J., Robinson J., Adiconis X., Fan L., Koziol M. J., Gnirke A., Nusbaum C., et al., 2010. Ab initio reconstruction of cell type-specific transcriptomes in mouse reveals the conserved multi-exonic structure of lincRNAs. Nature Biotechnology, 28(5):503–510.

39. Guttman M. and Rinn J. L., 2012. Modular regulatory principles of large non-coding RNAs. Nature, 482(7385):339–346.

40. Hansen K. D., Irizarry R. A., and Zhijin W., 2012. Removing technical variability in RNA-seq data using conditional quantile normalization. Biostatistics, 13(2):204–216.

41. Heid I. M., Jackson A. U., Randall J. C., Winkler T. W., Qi L., Steinthorsdottir V., Thorleifs-son G., Zillikens M. C., Speliotes E. K., Mägi R., et al., 2010. Meta-analysis identifies 13 new loci associated with waist-hip ratio and reveals sexual dimorphism in the genetic basis of fat distribution. Nature Genetics, 42(11):949–960.

42. Herriges M. J., Swarr D. T., Morley M. P., Rathi K. S., Peng T., Stewart K. M., and Morrisey E. E., 2014. Long noncoding rnas are spatially correlated with transcription factors and regulate lung development. Genes Dev., 28(12):1363–1379.

43. Hindorff L. A., Sethupathy P., Junkins H. A., Ramos E. M., Mehta J. P., Collins F. S., and Manolio T. A., 2009. Potential etiologic and functional implications of genome-wide association loci for human diseases and traits. Proceedings of the National Academy of Sciences, 106(23):9362–7.

44. H¨oglinger G. U., Melhem N. M., Dickson D. W., Sleiman P. M., Wang L.-S., Klei L., Rade-makers R., de Silva R., Litvan I., Riley D. E., et al., 2011. Identification of common variants influencing risk of the tauopathy progressive supranuclear palsy. Nature Genetics, 43(7):699–705.

45. Houlston R. S., Cheadle J., Dobbins S. E., Tenesa A., Jones A. M., Howarth K., Spain S. L., Broderick P., Domingo E., Farrington S., et al., 2010. Meta-analysis of three genome-wide association studies identifies susceptibility loci for colorectal cancer at 1q41, 3q26. 2, 12q13. 13 and 20q13. 33. Nature Genetics, 42(11):973–977.

46. Huarte M., Guttman M., Feldser D., Garber M., Koziol M. J., Kenzelmann-Broz D., Khalil A. M., Zuk O., Amit I., Rabani M., et al., 2010. A large intergenic noncoding RNA induced by p53 mediates global gene repression in the p53 response. Cell, 142(3):409–419.

47. Ishii N., Ozaki K., Sato H., Mizuno H., Saito S., Takahashi A., Miyamoto Y., Ikegawa S., Kamatani N., Hori M., et al., 2006. Identification of a novel non-coding RNA, *MIAT*, that confers risk of myocardial infarction. Journal of Human Genetics, 51(12):1087–1099.

48. Jeggari A., Marks D. S., and Larsson E., 2012. miRcode: a map of putative microRNA target sites in the long non-coding transcriptome. Bioinformatics, 28(15):2062–2063.

49. Jendrzejewski J., He H., Radomska H. S., Li W., Tomsic J., Liyanarachchi S., Davuluri R. V., Nagy R., and de la Chapelle, A., 2012. The polymorphism rs944289 predisposes to papillary thyroid carcinoma through a large intergenic noncoding RNA gene of tumor suppressor type. Proceedings of the National Academy of Sciences, 109(22):8646–8651.

50. Khalil A. M., Guttman M., Huarte M., Garber M., Raj A., Morales D. R., Thomas K., Presser A., Bernstein B. E., van Oudenaarden A., et al., 2009. Many human large intergenic noncoding RNAs associate with chromatin-modifying complexes and affect gene expression. Proceedings of the National Academy of Sciences, 106(28):11667–11672.

51. Klattenhoff C. A., Scheuermann J. C., Surface L. E., Bradley R. K., Fields P. A., Steinhauser M. L., Ding H., Butty V. L., Torrey L., Haas S., et al., 2013. Braveheart, a long noncoding rna required for cardiovascular lineage commitment. Cell, 152(3):570–583.

52. K¨ottgen A., Albrecht E., Teumer A., Vitart V., Krumsiek J., Hundertmark C., Pistis G., Ruggiero D., O’Seaghdha C. M., Haller T., et al., 2013. Genome-wide association analyses identify 18 new loci associated with serum urate concentrations. Nature Genetics, 45(2):145–154.

53. Kretz M., Siprashvili Z., Chu C., Webster D. E., Zehnder A., Qu K., Lee C. S., Flockhart R. J., Groff A. F., Chow J., et al., 2013. Control of somatic tissue differentiation by the long non-coding RNA TINCR. Nature, 493(7431):231–235.

54. Lewis B. P., Burge C. B., and Bartel D. P., 2005. Conserved seed pairing, often flanked by adenosines, indicates that thousands of human genes are microRNA targets. Cell, 120(1):15–20.

55. Liao Y., Smyth G. K., and Shi W., 2014. featureCounts: an efficient general purpose program for assigning sequence reads to genomic features. Bioinformatics, 30(7):923–930.

56. Lindgren C. M., Heid I. M., Randall J. C., Lamina C., Steinthorsdottir V., Qi L., Speliotes E. K., Thorleifsson G., Willer C. J., Herrera B. M., et al., 2009. Genome-wide association scan meta-analysis identifies three loci influencing adiposity and fat distribution. PLoS Genetics, 5(6):e1000508.

57. Loewer S., Cabili M. N., Guttman M., Loh Y.-H., Thomas K., Park I. H., Garber M., Curran M., Onder T., Agarwal S., et al., 2010. Large intergenic non-coding rna-ror modulates reprogramming of human induced pluripotent stem cells. Nature Genetics, 42(12):1113–1117.

58. Lonsdale J., Thomas J., Salvatore M., Phillips R., Lo E., Shad S., Hasz R., Walters G., Garcia F., Young N., et al., 2013. The genotype-tissue expression (GTEx) project. Nature Genetics, 45(6):580–585.

59. Lorenz R., Bernhart S. H., Zu Siederdissen, C. H., Tafer H., Flamm C., Stadler P. F., Hofacker I. L., et al., 2011. ViennaRNA Package 2.0. Algorithms for Molecular Biology, 6(1):26.

60. Mangravite L. M., Engelhardt B. E., Medina M. W., Smith J. D., Brown C. D., Chasman D. I., Mecham B. H., Howie B., Shim H., Naidoo D., et al., 2013. A statin-dependent QTL for GATM expression is associated with statin-induced myopathy. Nature, 502(7471):377–80.

61. Manning A. K., Hivert M.-F., Scott R. A., Grimsby J. L., Bouatia-Naji N., Chen H., Rybin D., Liu C.-T., Bielak L. F., Prokopenko I., et al., 2012. A genome-wide approach accounting for body mass index identifies genetic variants influencing fasting glycemic traits and insulin resistance. Nature Genetics, 44(6):659–669.

62. Manolio T. A., Collins F. S., Cox N. J., Goldstein D. B., Hindorff L. A., Hunter D. J., McCarthy M. I., Ramos E. M., Cardon L. R., Chakravarti A., et al., 2009. Finding the missing heritability of complex diseases. Nature, 461(7265):747–753.

63. Mattick J. S., 2009. The genetic signatures of noncoding RNAs. PLoS Genetics, 5(4):e1000459.

64. McVicker G., van de Geijn, B., Degner J. F., Cain C. E., Banovich N. E., Raj A., Lewellen N., Myrthil M., Gilad Y., and Pritchard J. K., et al., 2013. Identification of genetic variants that affect histone modifications in human cells. Science, 342(6159):747–749.

65. Miyagawa R., Tano K., Mizuno R., Nakamura Y., Ijiri K., Rakwal R., Shibato J., Masuo Y., Mayeda A., Hirose T., et al., 2012. Identification of cis-and trans-acting factors involved in the localization of malat-1 noncoding rna to nuclear speckles. RNA, 18(4):738–751.

66. Montgomery S. B. and Dermitzakis E. T., 2011. From expression QTLs to personalized transcriptomics. Nature Reviews Genetics, 12(4):277–282.

67. Montgomery S. B., Sammeth M., Gutierrez-Arcelus M., Lach R. P., Ingle C., Nisbett J., Guigo R., and Dermitzakis E. T., 2010. Transcriptome genetics using second generation sequencing in a caucasian population. Nature, 464(7289):773–777.

68. Nakashima M., Chung S., Takahashi A., Kamatani N., Kawaguchi T., Tsunoda T., Hosono N., Kubo M., Nakamura Y., and Zembutsu H., et al., 2010. A genome-wide association study identifies four susceptibility loci for keloid in the japanese population. Nature Genetics, 42(9):768–771.

69. Nica A. C., Montgomery S. B., Dimas A. S., Stranger B. E., Beazley C., Barroso I., and Dermitzakis E. T., 2010. Candidate causal regulatory effects by integration of expression QTLs with complex trait genetic associations. PLoS Genetics, 6(4):e1000895.

70. Nica A. C., Parts L., Glass D., Nisbet J., Barrett A., Sekowska M., Travers M., Potter S., Grundberg E., Small K., et al., 2011. The architecture of gene regulatory variation across multiple human tissues: the muther study. PLoS Genetics, 7(2):e1002003.

71. Nicolae D. L., Gamazon E., Zhang W., Duan S., Dolan M. E., and Cox N. J., 2010. Trait-Associated SNPs Are More Likely to Be eQTLs: Annotation to Enhance Discovery from GWAS. PLoS Genetics, 6(4).

72. Nie L., Wu H.-J., Hsu J.-M., Chang S.-S., LaBaff A. M., Li C.-W., Wang Y., Hsu J. L., and Hung M.-C., 2012. Long non-coding RNAs: versatile master regulators of gene expression and crucial players in cancer. Am. J. Transl. Res., 4(2):127.

73. Ørom U. A., Derrien T., Beringer M., Gumireddy K., Gardini A., Bussotti G., Lai F., Zytnicki M., Notredame C., Huang Q., et al., 2010. Long noncoding RNAs with enhancer-like function in human cells. Cell, 143(1):46–58.

74. Pai A. A., Cain C. E., Mizrahi-Man O., De Leon S., Lewellen N., Veyrieras J.-B., Degner J. F., Gaffney D. J., Pickrell J. K., Stephens M., et al., 2012. The contribution of RNA decay quantitative trait loci to inter-individual variation in steady-state gene expression levels. PLoS Genetics, 8(10):e1003000.

75. Pedersen J. S., Bejerano G., Siepel A., Rosenbloom K., Lindblad-Toh K., Lander E. S., Kent J., Miller W., and Haussler D., 2006. Identification and classification of conserved RNA secondary structures in the human genome. PLoS Computational Biology, 2(4):e33.

76. Pedregosa F., Varoquaux G., Gramfort A., Michel V., Thirion B., Grisel O., Blondel M., Prettenhofer P., Weiss R., Dubourg V., et al., 2011. Scikit-learn: Machine learning in Python. The Journal of Machine Learning Research, 12:2825–2830.

77. Petersen T. N., Brunak S., von Heijne G., and Nielsen H., 2011. SignalP 4.0: discriminating signal peptides from transmembrane regions. Nature Methods, 8(10):785–786.

78. Petruk S., Sedkov Y., Riley K. M., Hodgson J., Schweisguth F., Hirose S., Jaynes J. B., Brock H. W., and Mazo A., 2006. Transcription of bxd noncoding RNAs promoted by trithorax represses Ubx in cis by transcriptional interference. Cell, 127(6):1209–1221.

79. Pickrell J. K., Marioni J. C., Pai A. A., Degner J. F., Engelhardt B. E., Nkadori E., Veyri-eras, J.-B., Stephens M., Gilad Y., and Pritchard J. K., et al., 2010. Understanding mechanisms underlying human gene expression variation with RNA sequencing. Nature, 464(7289):768–772.

80. Ponjavic J., Ponting C. P., and Lunter G., 2007. Functionality or transcriptional noise? Evidence for selection within long noncoding RNAs. Genome Research, 17(5):556–65.

81. Popadin K., Gutierrez-Arcelus M., Dermitzakis E. T., and Antonarakis S. E., 2013. Genetic and epigenetic regulation of human lincRNA gene expression. American Journal of Human Genetics, 93(6):1015–26.

82. Porcu E., Medici M., Pistis G., Volpato C. B., Wilson S. G., Cappola A. R., Bos S. D., Deelen J., den Heijer M., Freathy R. M., et al., 2013. A meta-analysis of thyroid-related traits reveals novel loci and gender-specific differences in the regulation of thyroid function. PLoS Genetics, 9(2):e1003266.

83. Prensner J. R., Iyer M. K., Sahu A., Asangani I. A., Cao Q., Patel L., Vergara I. A., Davicioni E., Erho N., Ghadessi M., et al., 2013. The long noncoding rna schlap1 promotes aggressive prostate cancer and antagonizes the swi/snf complex. Nature Genetics.

84. Price A. L., Helgason A., Thorleifsson G., McCarroll S. A., Kong A., and Stefansson K., 2011. Single-tissue and cross-tissue heritability of gene expression via identity-by-descent in related or unrelated individuals. PLoS Genetics, 7(2):e1001317.

85. Quek X. C., Thomson D. W., Maag J. L., Bartonicek N., Signal B., Clark M. B., Gloss B. S., and Dinger M. E., 2014. lncrnadb v2. 0: expanding the reference database for functional long noncoding rnas. Nucleic Acids Research gku988.

86. Randall J. C., Winkler T. W., Kutalik Z., Berndt S. I., Jackson A. U., Monda K. L., Kilpeläinen T. O., Esko T., Mägi R., Li S., et al., 2013. Sex-stratified genome-wide association studies including 270,000 individuals show sexual dimorphism in genetic loci for anthropometric traits. PLoS Genetics, 9(6):e1003500.

87. Rawal R., Teumer A., V¨olzke H., Wallaschofski H., Ittermann T., Åsvold, B. O., Bjoro T., Greiser K. H., Tiller D., Werdan K., et al., 2012. Meta-analysis of two genome-wide association studies identifies four genetic loci associated with thyroid function. Human Molecular Genetics, dds136.

88. Rinn J. L. and Chang H. Y., 2012. Genome regulation by long noncoding rnas. Annu. Rev. Biochem., 81.

89. Rinn J. L., Kertesz M., Wang J. K., Squazzo S. L., Xu X., Brugmann S. A., Goodnough L. H., Helms J. A., Farnham P. J., Segal E., et al., 2007. Functional Demarcation of Active and Silent Chromatin Domains in Human HOX Loci by Noncoding RNAs. Cell, 129(7):1311–1323.

90. Sati S., Ghosh S., Jain V., Scaria V., and Sengupta S., 2012. Genome-wide analysis reveals distinct patterns of epigenetic features in long non-coding RNA loci. Nucleic Acids Research, 40(20):10018–10031.

91. Sauvageau M., Goff L. A., Lodato S., Bonev B., Groff A. F., Gerhardinger C., Sanchez-Gomez, D.B., Hacisuleyman E., Li E., Spence M., et al., 2013. Multiple knockout mouse models reveal lincrnas are required for life and brain development. eLife, 2:e01749.

92. Servin B. and Stephens M., 2007. Imputation-based analysis of association studies: candidate regions and quantitative traits. PLoS Genetics, 3(7):e114.

93. Siepel A., Bejerano G., Pedersen J. S., Hinrichs A. S., Hou M., Rosenbloom K., Clawson H., Spieth J., Hillier L. W., Richards S., et al., 2005. Evolutionarily conserved elements in vertebrate, insect, worm, and yeast genomes. Genome Research, 15(8):1034–1050.

94. Smith G. D. and Ebrahim S., 2003. Mendelian Randomization: can genetic epidemiology contribute to understanding environmental determinants of disease? International Journal of Epidemiology, 32(1):1–22.

95. Sun L., Goff L. A., Trapnell C., Alexander R., Lo K. A., Hacisuleyman E., Sauvageau M., Tazon-Vega B., Kelley D. R., Hendrickson D. G., et al., 2013. Long noncoding RNAs regulate adipogenesis. Proceedings of the National Academy of Sciences, 110(9):3387–3392.

96. Teumer A., Rawal R., Homuth G., Ernst F., Heier M., Evert M., Dombrowski F., V¨olker U., Nauck M., Radke D., et al., 2011. Genome-wide association study identifies four genetic loci associated with thyroid volume and goiter risk. The American Journal of Human Genetics, 88(5):664–673.

97. Tsai M.-C., Manor O., Wan Y., Mosammaparast N., Wang J. K., Lan F., Shi Y., Segal E., and Chang H. Y., 2010. Long noncoding RNA as modular scaffold of histone modification complexes. Science, 329(5992):689–693.

98. Ulitsky I. and Bartel D. P., 2013. lincRNAs: genomics, evolution, and mechanisms. Cell, 154(1):26–46.

99. Ulitsky I., Shkumatava A., Jan C. H., Sive H., and Bartel D. P., 2011. Conserved function of lincRNAs in vertebrate embryonic development despite rapid sequence evolution. Cell, 147(7):1537–1550.

100. Veyrieras J.-B., Kudaravalli S., Kim S. Y., Dermitzakis E. T., Gilad Y., Stephens M., and Pritchard J. K., 2008. High-resolution mapping of expression-QTLs yields insight into human gene regulation. PLoS Genetics, 4(10):e1000214.

101. Wang L., Park H. J., Dasari S., Wang S., Kocher J.-P., and Li W., 2013. Cpat: Coding-potential assessment tool using an alignment-free logistic regression model. Nucleic Acids Research, 41(6):e74–e74.

102. Wapinski O. and Chang H. Y., 2011. Long noncoding RNAs and human disease. Trends Cell Biol., 21(6):354–361.

103. Welter D., MacArthur J., Morales J., Burdett T., Hall P., Junkins H., Klemm A., Flicek P., Manolio T., Hindorff L., et al., 2014. The NHGRI GWAS Catalog, a curated resource of SNP-trait associations. Nucleic Acids Research, 42(D1):D1001–D1006.

104. Westra H.-J., Peters M. J., Esko T., Yaghootkar H., Schurmann C., Kettunen J., Christiansen M. W., Fairfax B. P., Schramm K., Powell J. E., et al., 2013. Systematic identification of trans eQTLs as putative drivers of known disease associations. Nature Genetics, 45(10):1238–1243.

105. Zhao J., Ohsumi T. K., Kung J. T., Ogawa Y., Grau D. J., Sarma K., Song J. J., Kingston R. E., Borowsky M., and Lee J. T., et al., 2010. Genome-wide identification of polycomb-associated rnas by rip-seq. Molecular Cell, 40(6):939–953.

106. Zhao J., Sun B. K., Erwin J. A., Song J.-J., and Lee J. T., 2008. Polycomb proteins targeted by a short repeat RNA to the mouse × chromosome. Science, 322(5902):750–756.

107. Zhou Y., Zhong Y., Wang Y., Zhang X., Batista D. L., Gejman R., Ansell P. J., Zhao J., Weng C., and Klibanski A., et al., 2007. Activation of p53 by meg3 non-coding rna. Journal of Biological Chemistry, 282(34):24731–24742.

